# RFX6 at locus 6q22 confers metastasis and drug resistance in prostate cancer

**DOI:** 10.1101/2024.01.08.574758

**Authors:** Mengjie Zhong, Wenjie Xu, Pan Tian, Qin Zhang, Zixian Wang, Limiao Liang, Qixiang Zhang, Yuehong Yang, Ying Lu, Gong-Hong Wei

## Abstract

Genetic and nonmutational epigenetic alterations are cancer hallmark characteristics. However, the role of inherited cancer predisposition alleles in co-opting lineage factor epigenetic reprogramming and contributing to tumor progression remains elusive. Here the FinnGen cohort phenome-wide analysis, along with recent multiple genome-wide association studies, has consistently identified the rs339331-RFX6/6q22 locus associated with prostate cancer (PCa) risk across diverse populations. We uncover that rs339331 resides at a reprogrammed androgen receptor (AR) binding site in PCa tumors, with the T risk allele enhancing AR chromatin occupancy under androgen signaling. We establish that RFX6 is an AR-regulated gene, intricately linked with rs339331, exhibiting synergistic prognostic value for PCa recurrence and metastasis. Through comprehensive *in vitro* and *in vivo* studies, we establish the oncogenic functions of RFX6 in promoting PCa cell proliferation and metastasis. Mechanistically, RFX6 upregulates transcription factor HOXA10 that profoundly correlates with adverse PCa outcomes and is pivotal in RFX6-mediated PCa progression, facilitating the epithelial-mesenchymal transition (EMT) process and modulating the TGFβ/SMAD signaling axis. Clinically, HOXA10 elevation is associated with increased EMT scores, tumor advancement and PCa recurrence. Remarkably, reducing RFX6 expression restores responsiveness of enzalutamide-resistant PCa cells and tumors to treatment. Our study highlights an interplay of disrupted genetic and epigenetic mechanisms converging on prostate lineage AR signaling, resulting in abnormal expression of RFX6 conferring PCa pathogenesis and enzalutamide resistance.

## Introduction

Prostate cancer (PCa) is the second most common cancer and the fifth leading cause of cancer-related mortality in men globally^1^. In 2023, PCa accounts for 29% of all new cancer cases in men, ranking highest in male cases with a significant mortality rate^2^. PCa incidence shows notable regional and ethnic disparities, being more prevalent in Europe, North America, and South Africa, and less so in Asia and North Africa^1^. However, the incidence rate of PCa increased rapidly in China with an annual rise of 12.6% since 2000^3, 4^. While the five-year survival rate for localized PCa is near 100%, this drops to about 30% for metastatic cases^2^. Most PCa cases are indolent and may not require immediate intervention, yet roughly 10-20% progress to an aggressive, potentially fatal metastatic castration-resistant form (CRPC)^3, 5–7^. This underscores an urgent need for a deeper understanding of PCa progression and the genetic underpinnings of therapy resistance.

PCa critically depends on the prostate lineage factor androgen receptor (AR), making androgen deprivation therapy (ADT) the primary treatment that targets androgen signaling^8^. While initially effective, ADT often leads to most PCa patients eventually developing CRPC, a stage at which treatment options become significantly limited. Despite FDA approval of next-generation androgen receptor (AR)-targeted agents like abiraterone and enzalutamide for CRPC, resistance is inevitable in most cases, severely constraining therapeutic choices and prognosis^5–7, 9^. Studies have indicated that resistance to enzalutamide may be linked to AR genomic mutations^10, 11^, splicing variations observed in CRPC patients^12, 13^, and alterations in critical signaling pathways^14–16^. Moreover, androgen deprivation-induced epithelial-mesenchymal transition (EMT) and cellular plasticity may alter gene expression, diminishing tumor sensitivity to enzalutamide^17^. These challenges underscore an urgent need to develop advanced AR-targeted therapies to boost effectiveness and counter drug resistance.

RFX6, a member of the regulatory factor X family, is characterized by its unique winged helical DNA-binding domain^18^. It plays a crucial role in enteroendocrine differentiation and pancreatic development, particularly in the regulation of insulin-producing cells^19–22^. A seminal study highlighted its involvement in a complex genetic network that reduces insulin secretion by β cells^23^. Intriguingly, recent extensive genome-wide association studies (GWAS) across various ethnic groups have linked the RFX6/6q22 locus to an increased risk and progression of PCa^24–30^. These studies collectively analyzed data from 221,340 East Asian, 378,708 European, and 6,930 African American men, respectively **(Supplementary Table 1)**. Our previous study reported that mechanistically, this association seems to be mediated by the rs339331/6q22 enhancer and HOXB13^31^, while the prostate-lineage-specific AR signaling may also play roles in transforming the regulatory effects of this locus and facilitate the elevated expression of RFX6. An insightful study highlighted a direct role of rs339331 in regulating RFX6 expression and the involvement of androgen signaling in modulating the effects of rs339331/6q22 on PCa risk^32^. Despite these advancements, yet whether specific interactions between the inherited rs339331/6q22 allele and prostate-lineage AR signaling in PCa progression remain uncertain. Additionally, the biological functions of RFX6 in PCa are not well-understood, partly due to the scarcity of in vivo evidence. Considering the extensive reprogramming of AR cistromes observed in PCa development, as highlighted in several groundbreaking studies^33–38^, we hypothesize that AR signaling is critical in gene regulatory mechanisms underpinning PCa susceptibility, which is likely instrumental in linking the rs339331/RFX6/6q22 locus with PCa risk and progression.

This study hence endeavors to elucidate the role of RFX6 in PCa progression, unravel its molecular mechanisms, and explore the clinical implications of these insights. We found a strong link between RFX6 and aggressive PCa features and elucidated how the 6q22/RFX6 locus contributes to PCa severity. This includes a novel association between RFX6 and the expression of HOXA10 and TGFβ2. Our study particularly highlights an involvement of RFX6 in epithelial-mesenchymal transition (EMT), primarily by upregulating HOXA10, a potential oncogenic transcription factor with prognostic value in PCa. We provide direct evidence of RFX6 in regulating HOXA10, which subsequently alters TGFβ2 expression, impacting PCa metastasis. Crucially, our research also identified RFX6 as a factor responsive to AR signaling, contributing to enzalutamide resistance. This finding positions RFX6 as a potential marker for risk stratification and therapeutic targeting in PCa treatment.

## Result

### AR signaling is strongly involved in gene regulatory control at the RFX6/6q22 locus

The interplay among chromatin, gene regulatory elements, and transcription factors is crucial in gene expression regulation^39^. Single nucleotide polymorphism (SNP) variants identified via GWAS are key in modulating tumor-specific cis-regulatory elements^40^. In a comprehensive Phenome-Wide Association Analysis (PheWAS) using data from the FinnGen cohort (n = 377,277)^41^, rs339331 at the RFX6/6q22 locus demonstrated the highest association with PCa among 2272 disease endpoints **(Fig. 1a)**. Our previous research identified the rs339331 variant at a conserved, functional regulatory element, characterized by the binding of transcription factor HOXB13, FOXA1, and AR in the PCa cell line^31^. Further analysis using human tissue ChIP-seq data^34^ corroborated the extensive recruitment of AR at the rs339331 enhancer unique in tumors **(Fig. 1b)**, indicating that the germline variant rs339331 converges on an epigenetically reprogrammed AR binding site over human prostate tumorigenesis. This convergence may lead to increased RFX6 expression, facilitating PCa progression, as supported by our subsequent studies demonstrating a strong correlation between the PCa risk-associated RFX6/6q22 locus, AR activity, and androgen signaling.

**Figure 1.**
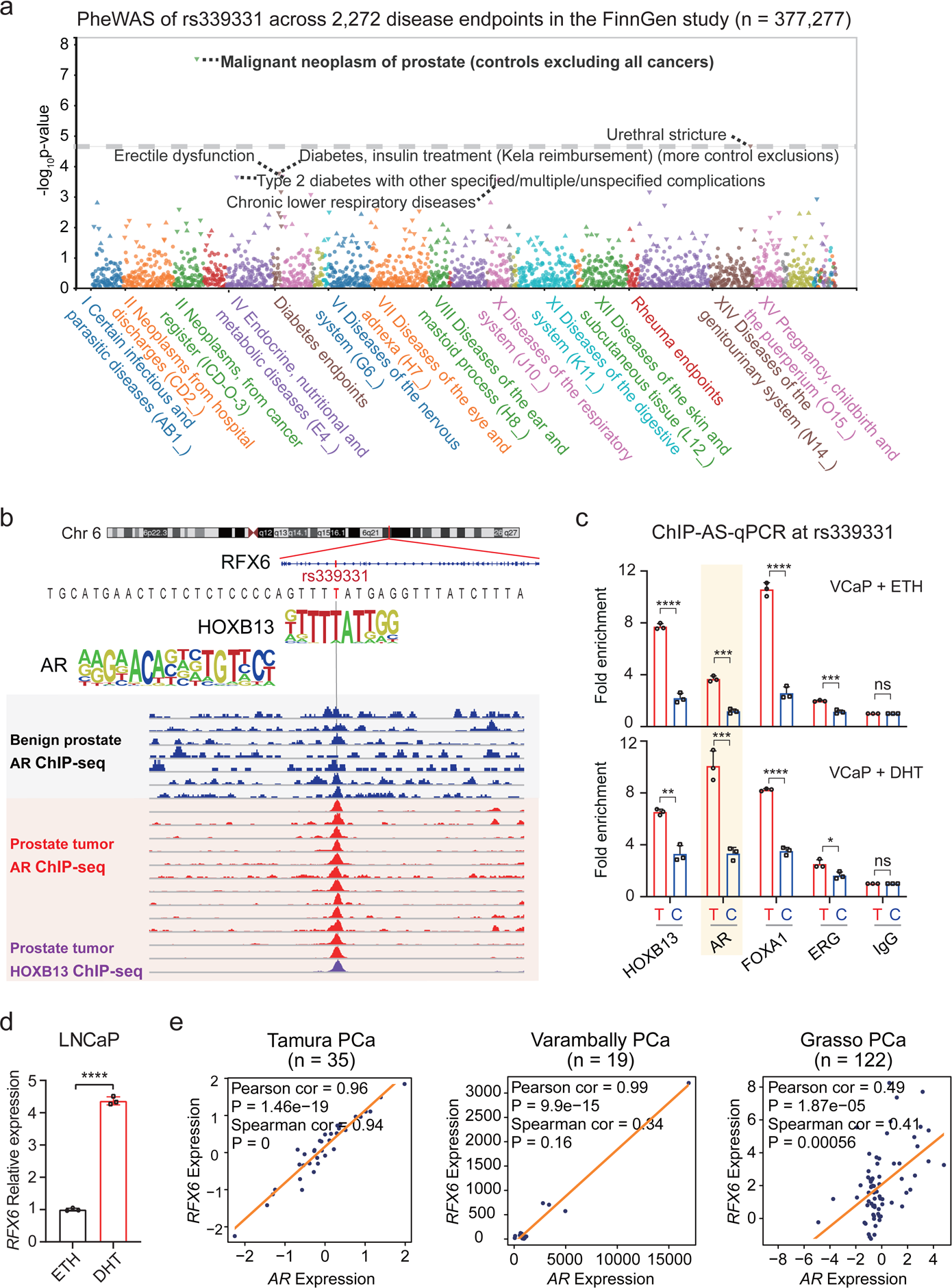
AR co-option of androgen signaling regulate RFX6 at the PCa risk-associated rs339331/6q22 locus. **a**, PheWAS outcomes from the FinnGen study (n = 377,277), showcasing associations between rs339331 and 2272 disease endpoints in. Associations are presented with P-values on the −log10 scale (vertical axis) against categories of disease endpoints (horizontal axis). The significance threshold was set at P = 3.28 × 10^−8^ (Bonferroni corrected). **b**, Genome browser views displaying AR ChIP-seq enrichment in prostate tumor tissues versus normal prostates at the rs339331/6q22 enhancer. **c**, ChIP followed by allele-specific (AS)-qPCR illustrating AR and other transcription factors preferentially binding to the T risk allele at rs339331, notably enhanced by DHT treatment. **d**, Quantitative real-time PCR analysis of RFX6 mRNA levels relative to GAPDH in LNCaP cells treated with DHT or ETH for 24 hours. **e**, Scatter plots demonstrating positive correlations between RFX6 and AR expression in prostate specimens from Tamura PCa (n = 35), Varambally PCa (n = 19) and Grasso PCa (n = 122) cohorts. In **c-d**, statistical significance assessed using a two-tailed Student’s t test. *p < 0.05, **p < 0.01, ***p < 0.001, ****p < 0.0001. In **e**, *p* values examined by the Pearson and Spearman correlation analysis.

Considering the pivotal role of AR in PCa, especially its notable enrichment at the rs339331 locus, we delved into its interplay with RFX6^42, 43^. Utilizing chromatin immunoprecipitation allele-specific quantitative PCR (ChIP-AS-qPCR) in VCaP cells, we confirmed significant AR recruitment to the PCa risk-associated T allele of rs339331 upon exposure to dihydrotestosterone (DHT) **(Fig. 1c).** We then examined the effect of androgen stimulation on RFX6 expression and found increased RFX6 levels in DHT-treated cells compared to controls **(Fig. 1d)**. Clinically, a robust correlation between RFX6 and AR expression was evident across multiple independent PCa cohorts^44–50^ **(Fig. 1e** and **Extended Data Fig. 1)**, corroborating the notion of androgen signaling induced RFX6 upregulation in a clinical context.

### Synergistic impact of rs339331 genotypes and RFX6 expression on PCa Severity

We further examined the relationship between RFX6 expression and the rs339331 genetic variation through an expression quantitative trait loci (eQTL) analysis using TCGA PCa cohort data^51^. This analysis demonstrated a significant association, with the T risk allele of rs339331 linked to increased RFX6 expression levels **(Fig. 2a)**, aligning with our prior findings in a Swedish cohort^31^. We then assessed the clinical relevance of RFX6 in PCa severity, employing prognostic evaluations in several independent PCa cohorts^46, 52, 53^. We observed a significant correlation between RFX6 upregulation and higher PCa grade or metastatic progression **(Fig. 2b-d)**. Elevated RFX6 expression was also noted in prostate tumors relative to paired normal tissues **(Fig. 2e)**. In mouse models with prostate-specific PTEN deletion, RFX6 expression markedly increased in tumor tissues compared to normal prostate **(Fig. 2f)**. Moreover, high RFX6 levels were linked to shorter progression-free survival in PCa patients^53^ **(Fig. 2g)**, corroborating its association with biochemical relapse^31, 54^, thereby highlighting the strong connection between RFX6 expression and PCa clinical severity.

**Figure 2.**
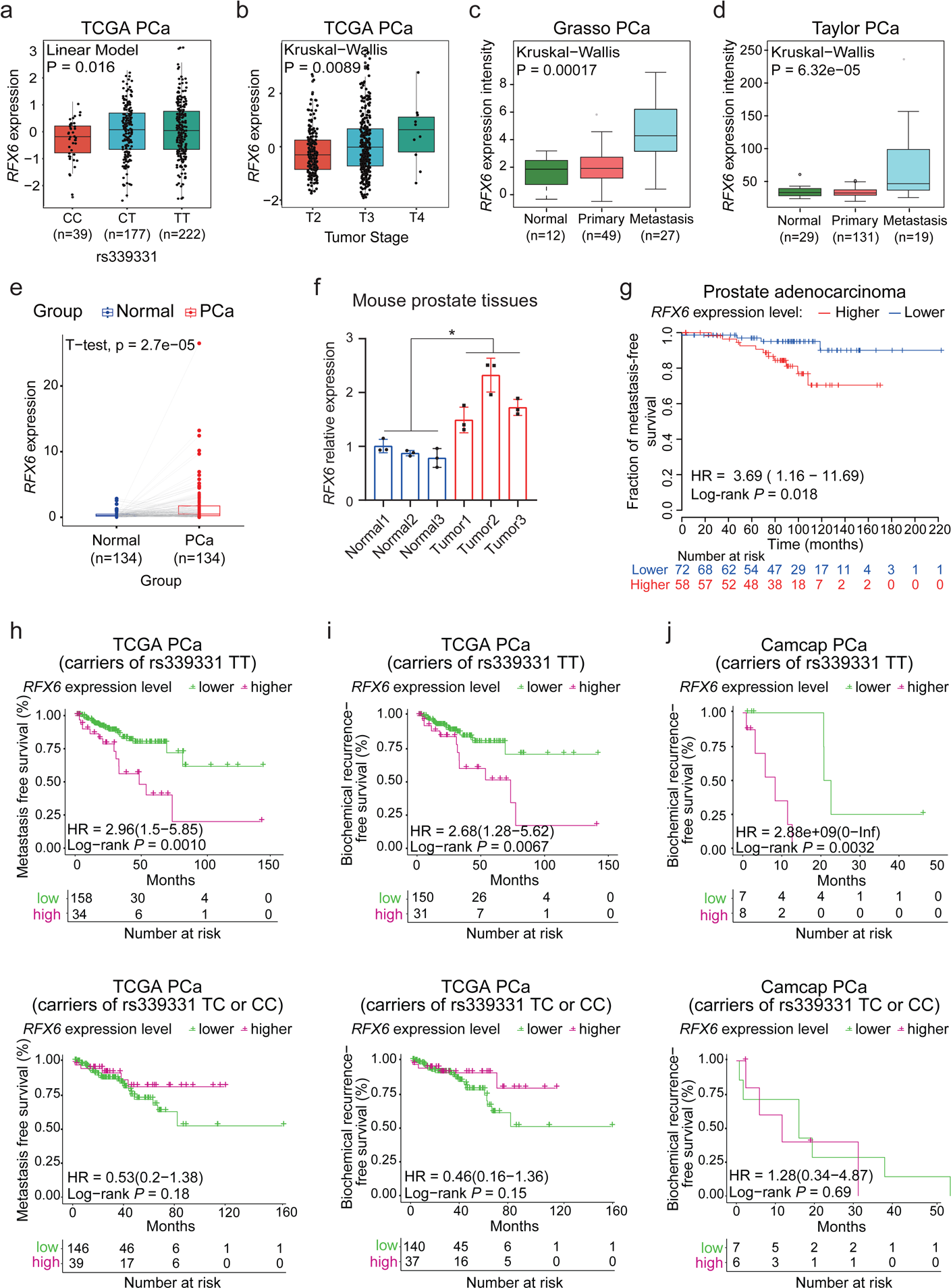
Association of elevated RFX6 expression with adverse clinical outcomes in PCa, linked to rs339331 TT risk genotype. **a**, Significant association of the PCa risk-associated rs339331 T allele with increased RFX6 expression. **b-d**, Analysis of clinical data demonstrating a correlation between elevated RFX6 expression and higher tumor stages (**b**) and progression to metastasis in PCa (**c,d**). **e**, Differential expression analysis of RFX6 in tumor-normal paired prostate specimens from a patient cohort. **f**, Analysis of relative RFX6 expression normalized to GAPDH in mouse tumors (PTEN prostate-specific knockout background) compared to normal murine prostate, using quantitative real-time PCR. **g**, Kaplan–Meier survival curve indicating worse metastasis outcomes in PCa patients with tumors exhibiting high RFX6 expression. **h-j**, RFX6 shows strong predictive value for PCa progression in patients with the rs339331 TT genotype (top panel), but not in patients with rs339331 CC or CT genotypes (bottom panel). In **a**, *p* values examined by linear model. In **b-d**, *p* values assessed by the Kruskal-Wallis tests. In **e-f**, statistical significance determined using a two-tailed Student’s t test. *p < 0.05, **p < 0.01, ***p < 0.001, ****p < 0.0001. In **g-j**, log-rank tests were applied for *p* values assessment.

Given the correlation of the rs339331 risk allele with increased PCa risk and elevated RFX6 expression, and association of RFX6 upregulation with PCa clinical severity, we examined the relationship between RFX6 expression and clinical indicators of tumor aggressiveness among PCa patients with varying rs339331 genotypes^52, 55^. We found that elevated RFX6 mRNA levels significantly correlate with shorter metastasis-free survival and a higher risk of biochemical recurrence, particularly in patients with the rs339331 TT genotype **(Fig. 2h-j)**. These findings highlight the pronounced upregulation of RFX6 expression associated with the TT/rs339331 variant in PCa tissues, and its overexpression, particularly in synergy with the T risk genotype of rs339331, is strongly linked to a poorer prognosis in PCa patients.

### RFX6 contributes to PCa cell metastasis potential

AR and androgen signaling are recognized as pivotal contributors to PCa pathogenesis^34, 43, 56, 57^ and RFX6 expression at the PCa risk 6q22 locus. We next explored the AR-dependent oncogenic potential of RFX6 in PCa development and progression. Initially, we assessed the effects of RFX6 on cellular proliferation and migration with androgen-sensitive PCa cells. Lentivirus-mediated RFX6 knockdown in 22Rv1 cells resulted in a notable decrease in cell growth, viability, colony formation, and migratory and invasive capabilities compared to control shRNA-treated cells **(Extended Data Fig. 2b-e)**, consistent with our previous findings using siRNA-mediated RFX6 tumor cell biology assays^31^.

To further substantiate these findings, we conducted a complementary approach by ectopic expression of RFX6 in 22Rv1 cells **(Fig. 3a** and **Extended Data Fig. 2a)**. This led to enhanced cell proliferation, migration, invasion, and clonal formation **(Fig. 3b-d)**. Additionally, flow cytometry analysis of 22Rv1 cells revealed that RFX6 overexpression increased the proportion of cells in the S phase and decreased those in the G1 phase, indicating a pro-proliferative effect **(Extended Data Fig. 2l)**. There was also a noticeable reduction in apoptosis in cells overexpressing RFX6 **(Extended Data Fig. 2m)**. Remarkably, enforced expression of RFX6 in AR-insensitive and RFX6-less-expressing PC3 and DU145 cells also significantly increased migration and invasion, underscoring the role of RFX6 in aggressive PCa cell behavior **(Extended Data Fig. 2f-k)**.

**Figure 3.**
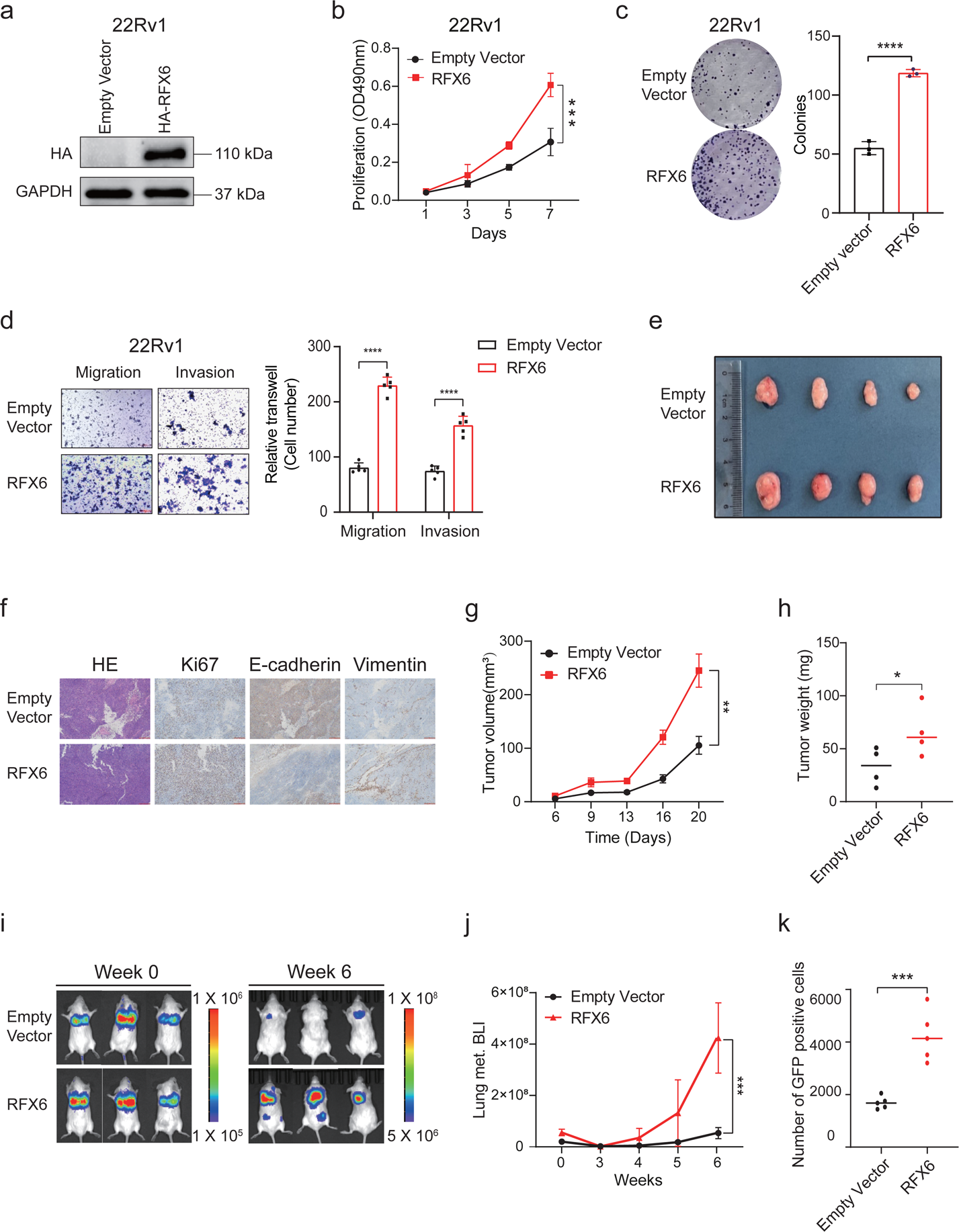
Impact of RFX6 upregulation on PCa tumor growth and metastasis *in vivo*. **a**, Immunoblots showcasing protein levels in 22Rv1 cells with RFX6 overexpression versus empty vector control. **b-c**, Analysis of cellular proliferation in 22Rv1 cells with RFX6 overexpression, measured by MTT assay (OD_490_; mean ± SD of three independent experiments) (**b**) and colony formation assay (**c**). **d**, Representative images from migration and invasion assays of 22Rv1 cells transfected with either empty vector or RFX6 expression construct. **e**, Images of excised xenograft tumors from mice groups with either empty vector or RFX6 overexpression. **f**, Hematoxylin and eosin (H&E) and IHC staining for Ki67, E-cadherin and vimentin in xenograft tumors overexpressing RFX6 versus empty vector groups. **g-h**, Quantification of tumor volumes (**g**) and masses (**h**) in excised xenograft tumors. **i**, In vivo imaging system (IVIS) images of intravenous 22Rv1 xenograft tumors at weeks 0 (left) and 6 (right) post-inoculation. Heatmap indicates IVIS signal intensity. **j**, Comparison of fluorescent intensity at weeks 0-6 post-injection, highlighting a significant increase in the RFX6 overexpression group compared to the empty vector group after 4 weeks. **k**, Flow cytometry (FACS) analysis of circulating GFP-positive 22Rv1 cell variants (CSCs) in the blood of SCID mice, with a scatter plot representing CSC counts per sample type (mean ± SD). Statistical Analysis: All data points were evaluated for statistical significance using two-tailed Student’s t-tests. *p < 0.05, **p < 0.01, ***p < 0.001, ****p < 0.0001.

Collectively, our results compellingly demonstrate the role of RFX6 in enhancing PCa cell proliferation and metastasis, applicable to both AR-positive and androgen-insensitive cell lines. This substantiates our earlier findings^31^ and expands our understanding of RFX6 functioning in PCa.

### RFX6 promotes PCa cellular proliferation and metastasis *in vivo*

To determine the effect of RFX6 on tumor growth *in vivo*, we conducted an experiment by subcutaneously implanting 22Rv1 cells, with either enforced RFX6 expression or an empty vector, into nude mice. The results showed that tumors derived from RFX6 overexpressing group exhibited significantly faster growth rates and markedly higher tumor weights compared to the control group **(Fig. 3e-h)**, clearly establishing its significant role in enhancing PCa tumor growth *in vivo*.

To assess the effect of RFX6 on metastatic potential *in vivo*, we developed lung metastasis model by intravenously injecting luciferase-labeled 22Rv1 cells with either elevated RFX6 expression or an empty vector. Six weeks after inoculation, *in vivo* imaging system (IVIS) revealed prominent metastatic foci in mice injected with RFX6 overexpressing cells, while the control group showed no significant IVIS signals **(Fig. 3i-j)**. Further, using flow cytometry sorting (FACS), we isolated GFP-positive cancer cells from the blood to analyze circulating cancer cells. This revealed a notably higher abundance of circulating cells with RFX6 expression compared to controls **(Fig. 3k)**, indicating the presence of circulating cells with enhanced survival capacity inside the bloodstream to potentiate their propensity for distant organs metastasis.

Taken together, these findings demonstrate that RFX6 not only promotes migration and invasion of PCa cells *in vitro* but also significantly contributes to the metastasis of transplanted tumors *in vivo*.

### RFX6 directly regulates HOXA10 expression

To understand how RFX6 influences the proliferation and metastasis capacities of PCa cells, we established a stable 22Rv1 cell line overexpressing RFX6 and performed RNA sequencing (RNA-seq) compared to control cells. Unexpectedly, the RNA-seq analysis indicated a modest impact, with only two genes HOXA10 and PPFIA4, showing altered expression patterns following RFX6 overexpression **(Fig. 4a)**. Further RT-qPCR validation confirmed consistent upregulation of HOXA10 by RFX6, while PPFIA4 did not exhibit similar consistency **(Fig. 4b** and **Extended Data Fig. 3a)**. Our studies also showed that RFX6 promotes HOXA10 expression at both transcriptional and protein levels **(Fig. 4b-c)**.

**Figure 4.**
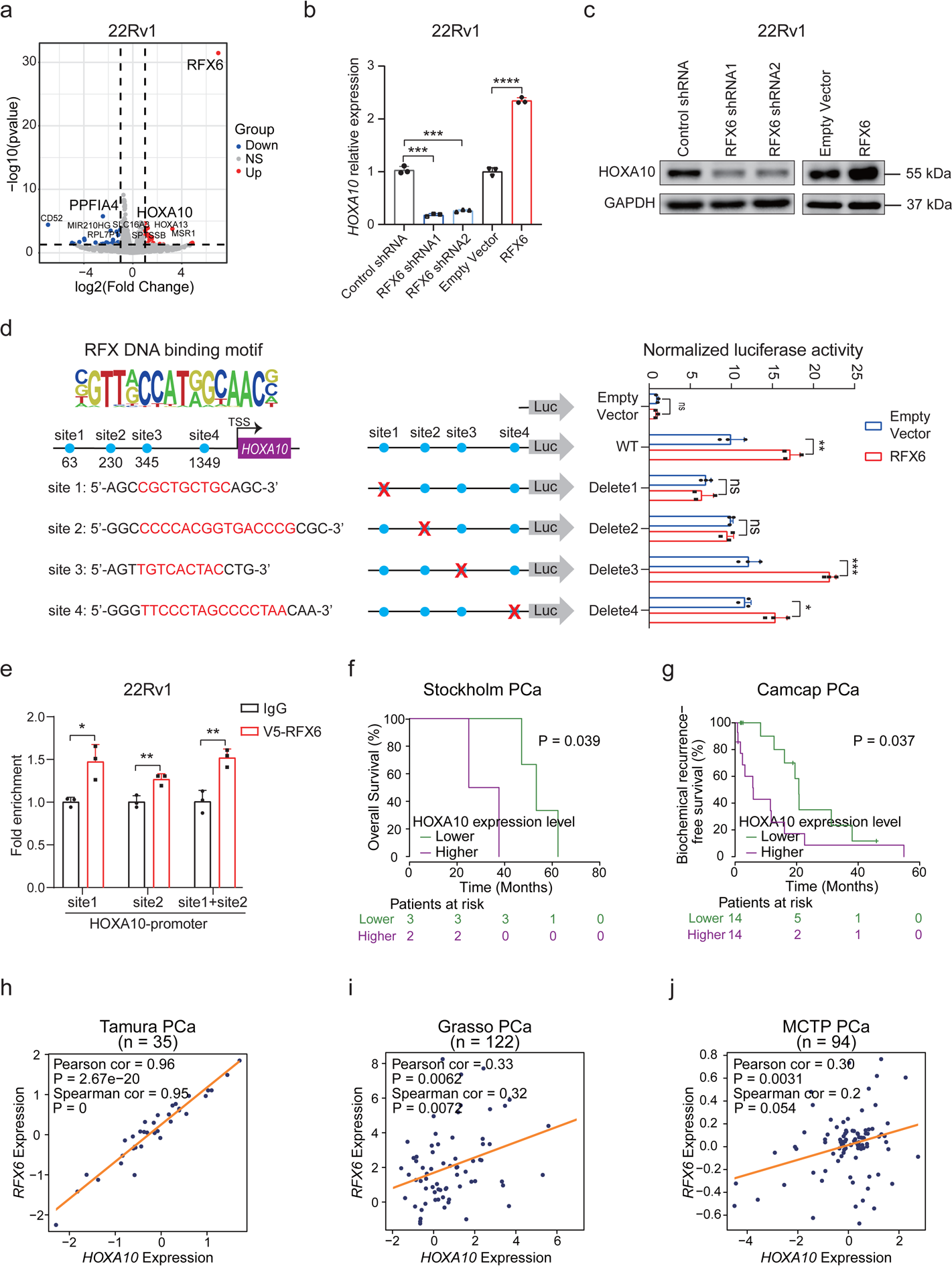
RFX6 directly regulates HOXA10 expression, correlating with poor prognosis in PCa. **a**, Volcano plot illustrating differential gene expression from RNA-seq in 22Rv1 cells with RFX6 overexpression vs. empty vector. Significant differentially expressed genes (DEGs; adjusted P-value < 0.05) are colored, with top-ranking DEGs highlighted. NS: not significant. **b**, Quantitative real-time PCR analysis of relative HOXA10 expression levels in 22Rv1 cells under indicated treatments. **c**, Immunoblots showing HOXA10 expression in cells with varying RFX6 expression statuses. **d**, Upper left: Predicted RFX DNA-binding motifs in HOXA10 promoter regions. Lower left: Schematic of wild-type/mutant HOXA10 promoter regions in the pGL4.10 luciferase reporter plasmid. Right: Luciferase activity quantification of HOXA10 promoter reporters in cells, with blue circles indicating RFX6-binding sites and red crosses denoting deleted sites. **e**, ChIP-qPCR confirmation of RFX6 binding at HOXA10 promoter sites 1 and 2 in 22Rv1 cells. **f-g**, Kaplan-Meier plots showing higher biochemical recurrence in PCa patients with elevated HOXA10 expression. **h-j**, Scatter plots revealing positive correlations between RFX6 and HOXA10 expression in prostate specimens from Tamura PCa (n = 35), Grasso PCa (n = 122) and MCTP PCa (n = 94) cohorts. In **b** and **d-e,** statistical significance assessed using the two-tailed Student’s t tests. *p < 0.05, **p < 0.01, ***p < 0.001, ****p < 0.0001. In **f-g**, *p* values were assessed using the log-rank tests, while Pearson and Spearman correlation analyses were conducted for panel **h-j**.

Given that RFX6 is a transcription factor, we examined its impact on HOXA10 promoter activity. The results indicated a significant increase of HOXA10 promoter transcriptional activity upon RFX6 transduction compared to the empty vector **(Extended Data Fig. 3b)**. We further identified and tested four potential RFX6 binding sites within the HOXA10 promoter using a dual-luciferase reporter assay **(Fig. 4d)**. The transcriptional activity was notably reduced upon deletion of binding sites 1 or 2 **(Fig. 4d)**. Further ChIP-qPCR analysis confirmed the recruitment of RFX6 to these sites within the HOXA10 promoter, demonstrating a direct regulatory influence of RFX6 on HOXA10 **(Fig. 4e)**.

Given that RFX6 confers PCa susceptibility and progression and directly regulates HOXA10 expression, we investigated whether HOXA10 correlates with PCa severity. This analysis revealed that HOXA10 showed increased expression in PCa tissues compared to normal prostate, and higher HOXA10 levels were associated with poorer prognosis^47, 52, 55, 58, 59^ **(Fig. 4f-g** and **Extended Data Fig. 3c-j)**. Remarkably, a robust correlation between RFX6 and HOXA10 expression was further substantiated through an analysis of PCa clinical data^44, 46^ **(Fig. 4h-j)**, suggesting that this regulatory circuit of RFX6 on HOXA10 is likely to be causal in the clinical setting. Collectively, our mechanistic studies reveal that RFX6 directly regulates HOXA10 expression by binding to its promoter, highlighting HOXA10 as a key target of RFX6 for its oncogenic function in PCa.

### HOXA10 upregulation correlates with adverse clinical outcomes and promotes aggressive tumor cell behavior in PCa

HOXA10, a member of the HOX gene family, is known for its crucial role in embryonic development and tumor progression^60–63^. Its specific contributions to PCa progression, however, remain largely unexplored. Our study analyzed HOXA10 expression in various PCa patient cohorts^46, 47, 52, 58, 59^ and found a significant correlation between its upregulation and advanced stages of PCa, particularly metastasis, and elevated levels of prostate-specific antigen (PSA) **(Fig. 5a-b** and **Extended Data Fig. 3c-j)**, the latter is a golden-standard biomarker for PCa screening and clinical decision-making^64^. Furthermore, in mouse models with PTEN prostate-specific knockout, HOXA10 expression was notably higher compared to wild-type controls **(Fig. 5c-d)**. These results suggest that RFX6-driven HOXA10 upregulation is closely associated with adverse clinical outcomes in PCa.

**Figure 5.**
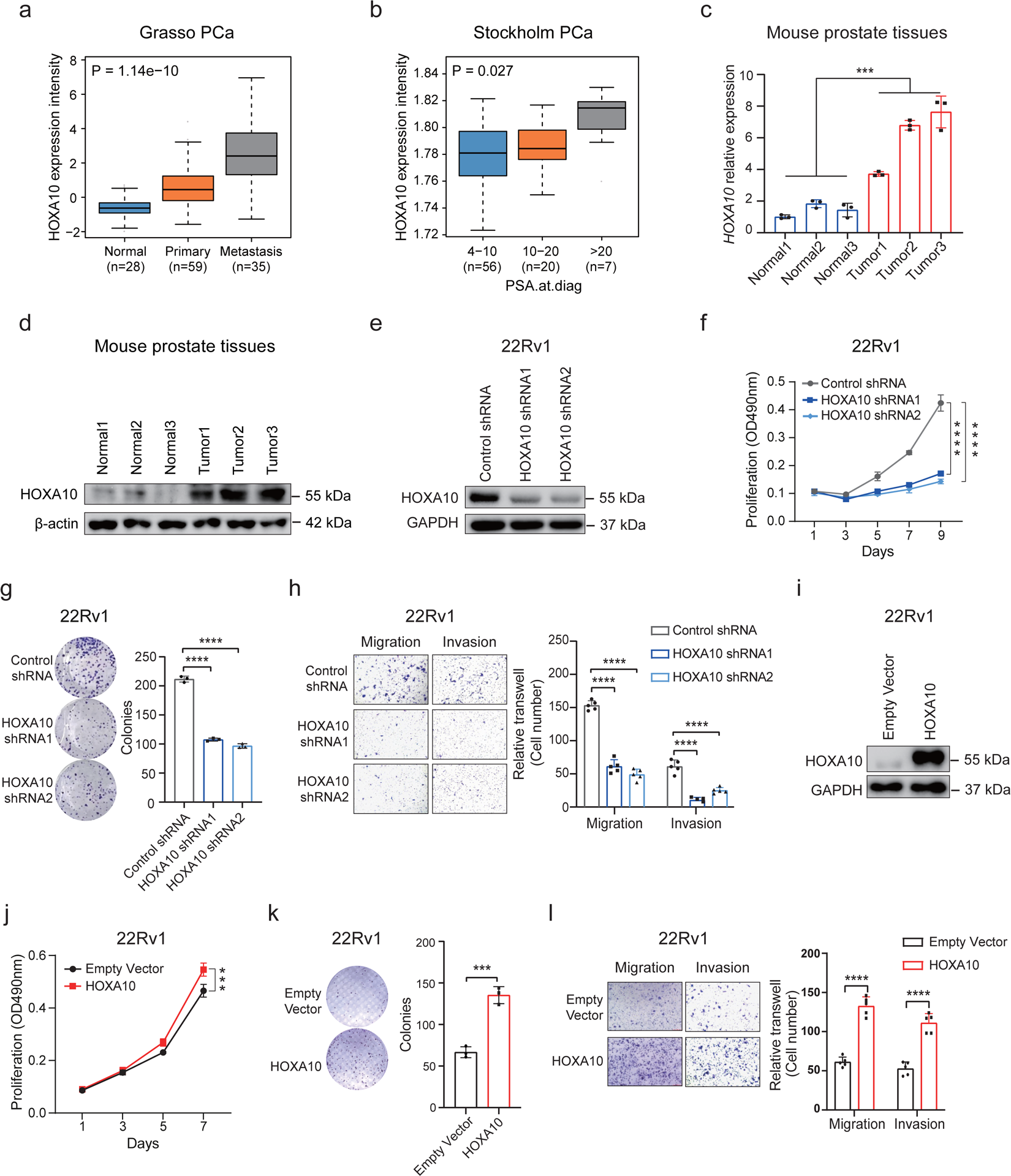
Evaluating the role of HOXA10 in PCa tumor cell phenotype. **a-b**, Clinical data analysis indicating a correlation between HOXA10 upregulation and tumor progression to metastasis (**a**) and increased levels of prostate-specific antigen (PSA) in PCa (**b**). **c-d**, Analysis of mRNA (**c**) and protein (**d**) expression levels of HOXA10 in mouse prostate tumors (prostate-specific PTEN knockout) compared to normal prostate tissue (wild type). **e**, Verification of the efficiency of shRNA-mediated HOXA10 knockdown through immunoblotting. **f-g**, Measurement of cellular proliferation in 22Rv1 cells with HOXA10 knockdown, assessed by MTT assay (OD_490_; mean ± SD from three independent experiments) (**f**) and colony formation assay (**g**). **h**, Representative images and quantification of migration and invasion in cells stably expressing shRNAs targeting HOXA10. **i**, Immunoblots showing HOXA10 expression in 22Rv1 cells with either empty vector or HOXA10 overexpression. **j-k**, Evaluation of cellular proliferation in 22Rv1 cells with enforced HOXA10 expression, using MTT assay (OD_490_; mean ± SD from three independent experiments) (**j**) and colony formation assay (**k**). **l**, Representative images of migration and invasion in 22Rv1 cells transfected with empty vector or HOXA10 expression construct. In **a-b**, *p* values examined by the Kruskal-Wallis tests. In **c** and **g-i** statistical significance assessed using the two-tailed Student’s t tests. Significance levels are indicated as *p < 0.05, **p < 0.01, ***p < 0.001, ****p < 0.0001.

To further understand the role of HOXA10 in PCa, we explored its effects on tumor cell phenotypes. Using shRNA-mediated knockdown models, we observed significant reduction in cell growth, viability, and colony formation in HOXA10-deficient cells compared to controls **(Fig. 5e-g)**. These cells also showed decreased migratory capability **(Fig. 5h** and **Extended Data Fig. 4a-c)**. Conversely, ectopic HOXA10 expression substantially potentiated cell proliferation, colony formation, and migration **(Fig. 5i-l** and **Extended Data Fig. 4d-f)**. These findings indicate that HOXA10 significantly influences cell proliferation, migration, invasion, and clonal formation, mirroring the tumor cellular phenotypic effects of RFX6 in PCa.

### HOXA10 plays critical roles in RFX6-driven PCa progression *in vitro* and *in vivo*

To examine whether the oncogenic effects of RFX6 on PCa cell proliferation and metastasis are phenotypically copied by HOXA10, we manipulated their expression in 22Rv1 cells **(Fig. 6a** and **Extended Data Fig. 5a)**. The results showed that HOXA10 depletion counteracted RFX6-induced increases in cell growth, proliferation, colony formation, migration, and resistance to apoptosis **(Extended Data Fig. 5b-e)**. In a subcutaneous xenograft model, RFX6 overexpression profoundly enhanced tumor growth, but this effect was significantly reduced with HOXA10 knockdown **(Fig. 6b-d)**. Immunohistochemistry (IHC) staining revealed that tumors with RFX6 overexpression displayed decreased E-cadherin staining and enhanced N-cadherin, vimentin, and Ki67 staining while these phenotypic changes were reversed when HOXA10 was knocked down **(Fig. 6e)**. Collectively, these findings confirm that HOXA10 mimics the effects of RFX6 on orchestrating aggressive PCa cellular behavior *in vitro* and tumor growth *in vivo*.

**Figure 6.**
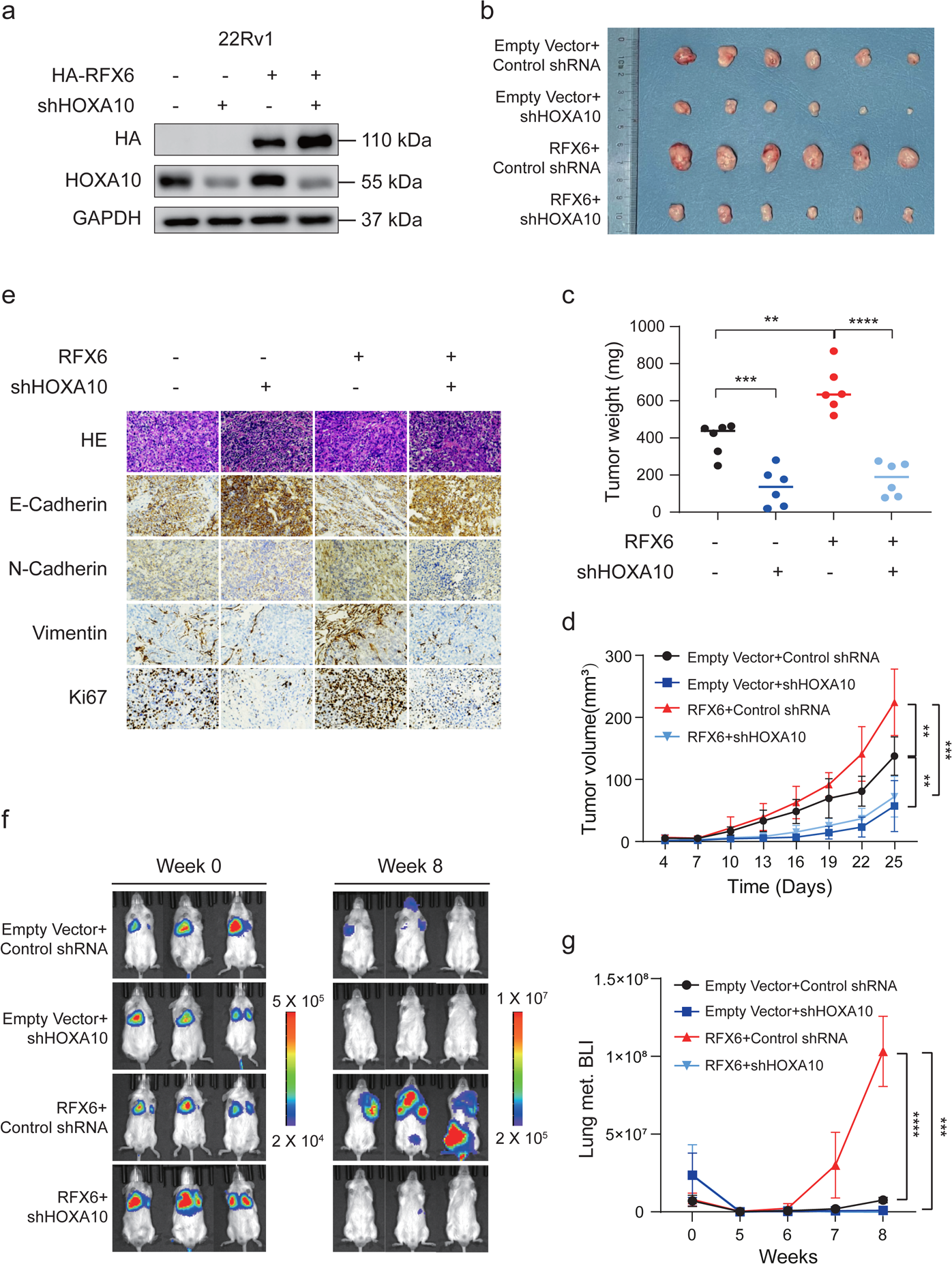
Phenocopying of RFX6 tumor-promoting effect by HOXA10 in *in vivo* proliferation and metastasis models. **a**, Immunoblotting analysis demonstrating the expression of RFX6 and HOXA10 in the cells. **b-d**, Visual representation of excised xenograft tumors (b), with quantification of tumor masses (**c**) and volumes (**d**) in the indicated experimental groups. **e**, Hematoxylin and eosin (H&E) and immunohistochemistry (IHC) staining for E-cadherin, N-cadherin, vimentin and Ki67 in xenograft tumors from the specified groups. **f**, In vivo imaging system (IVIS) imaging of intravenous 22Rv1 xenograft tumors in mice at weeks 0 (left) and 7 (right) post-inoculation, with a heatmap indicating signal intensity. **g**, Comparison of fluorescent intensity at 0-8 weeks post-injection, showing significantly increased fluorescence in the RFX6 overexpression groups after 6 weeks. Statistical Analysis: Statistical significance for all data was determined using two-tailed Student’s t-tests. Significance levels are indicated as *p < 0.05, **p < 0.01, ***p < 0.001, ****p < 0.0001.

We further evaluated the impact of HOXA10 on RFX6-driven PCa tumor progression using luciferase-labeled 22Rv1 cells in SCID mice. Overexpression of RFX6 resulted in enhanced metastasis by the seventh week, while HOXA10 downregulation led to reduced metastasis **(Fig. 6f-g)**. When RFX6 was upregulated and HOXA10 was simultaneously downregulated, the metastasis levels were similar to those observed with HOXA10 knockdown alone **(Fig. 6f-g)**. These patterns persisted into the eighth week **(Fig. 6g)**. Upon dissection, we noted significant lung metastasis in the RFX6 overexpression group, along with bone, brain metastasis, and lymph node enlargement, absent in other groups. In summary, while RFX6 overexpression consistently promoted metastasis **(Fig. 3i-j)**, its effect was significantly mitigated by HOXA10 knockdown, indicating that HOXA10 mimics the effect of RFX6 on metastatic progression in PCa.

### RFX6 propels PCa metastasis through HOXA10-induced EMT

Prior research has implicated the involvement of HOXA10 in epithelial-mesenchymal transition (EMT), a critical process in cancer metastasis^65,66^. In light of the significant changes in metastatic phenotypes following HOXA10 expression alteration observed in our study **(Fig. 6f-g)**, we sought to determine if these metastatic phenotypes are associated with EMT initiation driven by HOXA10 in PCa cells.

We thus assessed the correlation between HOXA10 expression and EMT scores in multiple PCa patient cohorts^48, 67, 68^, which revealed a consistent positive association **(Fig. 7a-c** and **Extended Data Fig. 6a-c)**. Using RT-qPCR and Western blotting, we investigated changes in key EMT markers in cell lines with altered HOXA10 expression. Overexpressing HOXA10 led to reduced levels of epithelial marker E-cadherin and increased mesenchymal markers such as N-cadherin, Vimentin, and EMT-associated transcription factors SNAIL1, TWIST1, and ZEB2 **(Fig. 7d** and **Extended Data Fig. 6d)**. Conversely, HOXA10 knockdown increased E-cadherin levels while reduced levels of mesenchymal markers and EMT-associated factors **(Fig. 7d** and **Extended Data Fig. 6d)**. Similar patterns were observed in cells with RFX6 overexpression or knockdown **(Extended Data Fig. 6e)**, highlighting the role of RFX6 in modulating HOXA10 expression and its impact on EMT marker expression.

**Figure 7.**
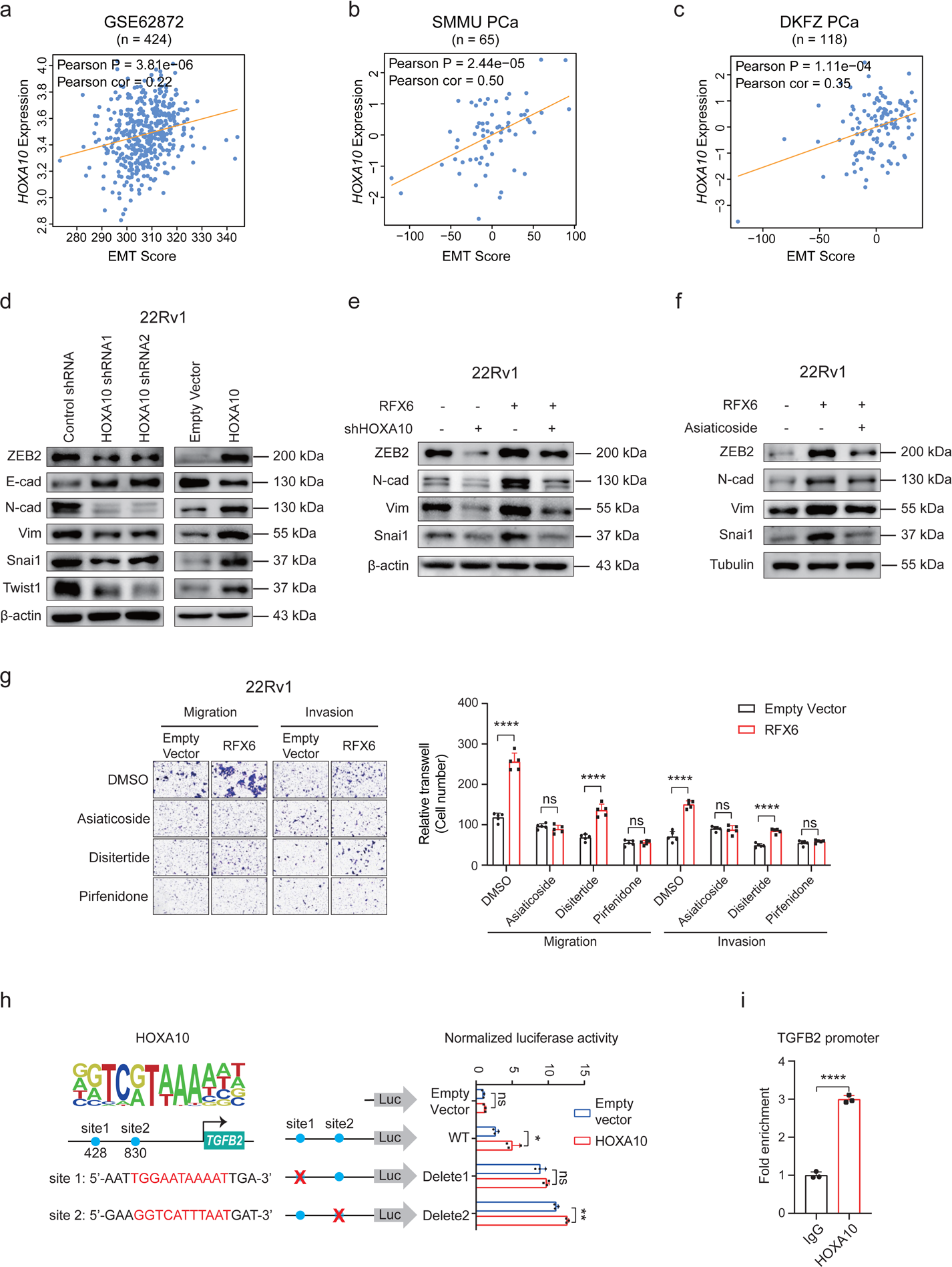
RFX6 mediates EMT via HOXA10-upregulated TGFβ/SMAD signaling. **a-c**, Correlation analysis showing a positive relationship between HOXA10 expression and EMT scores in three independent cohorts of PCa patients. **d-e**, Immunoblotting results depicting the expression of EMT-related molecules (E-cadherin, N-cadherin, Vimentin, Snail1, ZEB2, Twist1) in cells of the indicated groups. **f**, Analysis of EMT-associated molecules (N-cadherin, vimentin, Snail, ZEB2) in cells treated with Asiaticoside, assessed by immunoblotting. **g**, Changes in migratory and invasive capabilities in RFX6-overexpressing cells treated with Asiaticoside, Disitertide or Pirfenidone. **h**, Upper left: Predicted HOXA10 DNA-binding motifs in TGFβ2 promoter regions. Lower left: Schematic of wild-type/mutant TGFβ2 promoters in the pGL4.10 luciferase reporter plasmid. Right: Quantification of luciferase activity in cells with HOXA10-binding sites (blue circles) and deleted sites (red crosses). **i**, ChIP-qPCR verification of HOXA10 chromatin binding at the TGFβ2 promoter site 1 in 22Rv1 cells. Statistical Analysis: Pearson correlation analysis was used for p-value assessment in panels **a-c**. Two-tailed Student’s t-tests were employed for statistical significance in panels **g-i**, with significance levels indicated as *p < 0.05, **p < 0.01, ***p < 0.001, ****p < 0.0001.

In further experiments, we conducted HOXA10 knockdown in the context of RFX6 overexpression. This led to the reversal of E-cadherin downregulation typically seen with RFX6 overexpression. Additionally, HOXA10 knockdown mitigated the increased expression of N-cadherin, Vimentin, SNAIL1, and ZEB2 observed in the RFX6 overexpression group **(Fig. 7e)**. These results highlight a dynamic interplay among RFX6, HOXA10, and EMT markers, collectively playing a significant role in PCa progression.

### RFX6 influences TGFβ signaling in PCa

Given the known role of TGFβ signaling in driving EMT progression^69^, we investigated whether RFX6 fosters EMT and metastasis in PCa via this pathway. We thus treated 22Rv1 cells overexpressing RFX6 with the TGFβ/SMAD signaling inhibitor Asiaticoside, observing a reduction in ZEB2, N-cadherin, Vimentin and Snai1 expression levels **(Fig. 7f)**. This suggests RFX6-mediated EMT can be hindered by inhibiting the TGFβ/SMAD signaling pathway.

We further identified that RFX6 upregulation specifically increased TGFβ2 expression, with no notable impact on TGFβ1 or TGFβ3 **(Extended Data Fig. 6f)**. RFX6-induced cell migration and invasion were significantly curtailed by both Asiaticoside and Pirfenidone, a TGFβ2 inhibitor, but not by Disitertide, a TGFβ1 inhibitor) **(Fig. 7g)**. Western blotting confirmed a decrease in EMT marker levels when TGFβ2 activity was suppressed in RFX6 overexpressing cells using Pirfenidone **(Extended Data Fig. 6g)**. These findings collectively suggest that RFX6 modulates TGFβ signaling predominantly through TGFβ2 in PCa.

We next sought to explore the regulatory mechanisms underlying TGFβ2 expression in PCa. Using dual-luciferase reporter assays, we examined the transcriptional influence of RFX6 and HOXA10 on the TGFβ2 promoter. We found that while overexpression of HOXA10 significantly increased TGFβ2 promoter activity, RFX6 alone did not produce a similar effect **(Extended Data Fig. 6h-i)**. Through bioinformatics prediction, we identified potential HOXA10 binding sites within the TGFβ2 promoter. Notably, deletion of sites 1 abolished HOXA10-mediated modulation of TGFβ2 promoter activity, as verified by dual-luciferase reporter assays and ChIP-qPCR **(Fig. 7h-i)**. These findings indicate a unidirectional transcriptional regulatory relationship involving RFX6, HOXA10, and TGFβ2, with HOXA10 directly binding at site 1 of the TGFβ2 promoter.

### Inhibition of RFX6 curbs enzalutamide-resistant PCa proliferation *in vitro* and *in vivo*

Our investigations as described above have established a robust association between RFX6 expression and androgen signaling, alongside nonmutational epigenetically reprogrammed AR binding sites in tumors, with RFX6 showing higher expression in AR-dependent cell lines **(Fig. 1** and **Extended Data Fig. 1)**. In the clinical context, enzalutamide, an AR inhibitor, is widely used to treat CRPC. Notably, in our studies involving the CRPC cell line 22Rv1, treatment with enzalutamide resulted in an increased RFX6 expression in correlation with the drug concentration **(Fig. 8a)**.

**Figure 8.**
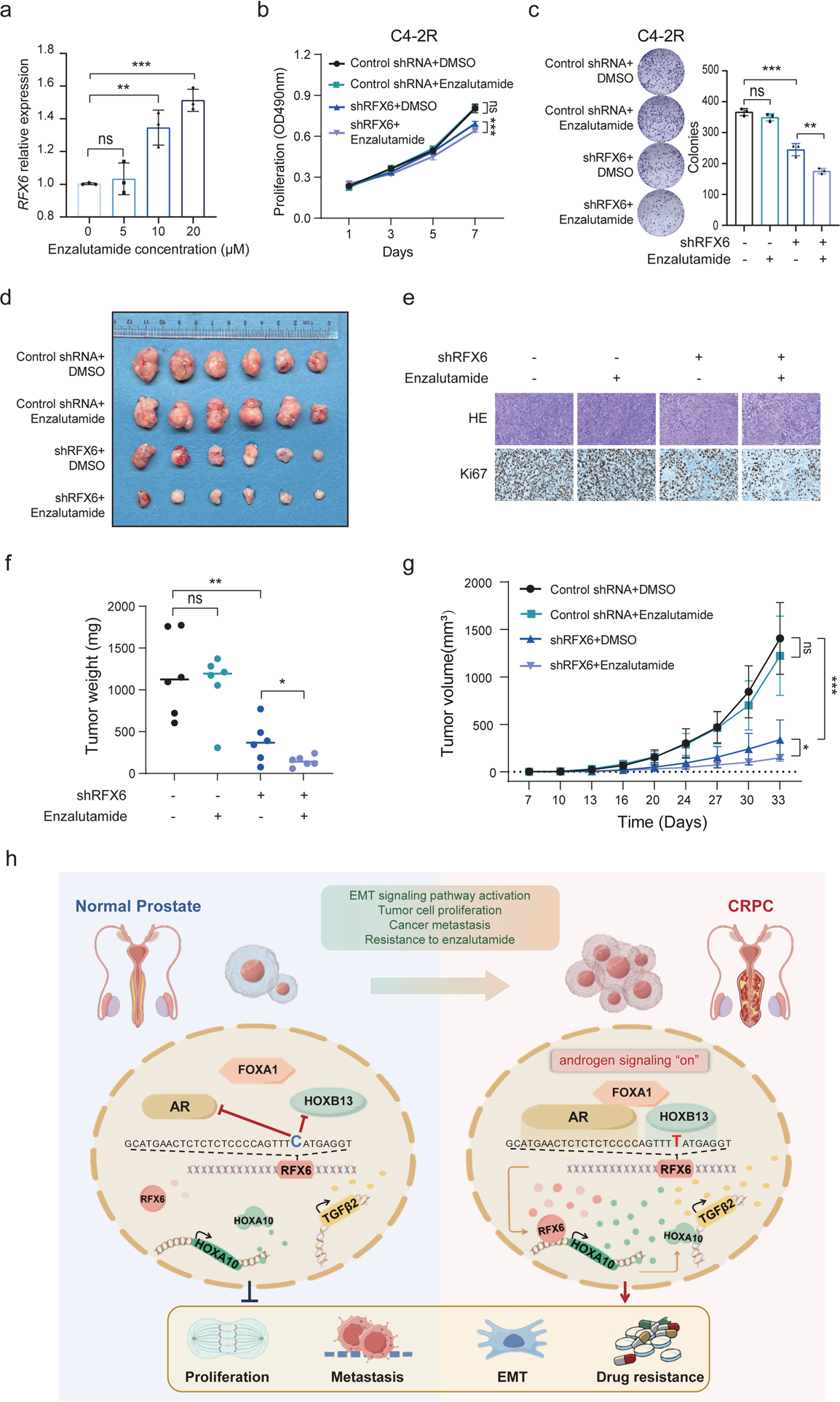
Enhanced tumor sensitivity to enzalutamide through diminished RFX6 expression. **a**, Real-time PCR analysis confirming RFX6 expression levels in 22Rv1 cells treated with varying concentrations of enzalutamide for 24 hours. **b-c**, The impact of RFX6 shRNA on the proliferation (**b**) and colony formation (**c**) of C4-2R cells, highlighting increased cell sensitivity to enzalutamide treatment. **d**, Images of excised xenograft tumors from different treatment groups. **e**, Hematoxylin and eosin (H&E) and immunohistochemistry (IHC) staining for Ki67 in xenograft tumors as indicated. **f-g**, Quantification of tumor masses (**f**) and volumes (**g**) in excised xenograft tumors from specified murine groups. **h**, Schematic model illustrating the regulatory pathway involving rs339331, RFX6, HOXA10, and TGFβ2. This pathway elucidates the mechanism by which AR binding at the rs339331/enhancer region of RFX6 gene leads to RFX6 upregulation in PCa tumors, subsequently increasing HOXA10 and TGFβ2 expression. These molecular events promote PCa cell proliferation, induce epithelial-mesenchymal transition (EMT), enhance metastasis, and contribute to enzalutamide resistance, culminating in the progression to castration-resistant PCa (CRPC). All the statistical significance assessed using the Two-tailed Student’s t tests, with significance levels indicated as *p < 0.05, **p < 0.01, ***p < 0.001, ****p < 0.0001.

Given the known link between high RFX6 expression and poor clinical prognosis in PCa, as well as its roles in disease progression, we investigated whether increased RFX6 expression may lead to resistance during enzalutamide treatment. We modulated RFX6 expression in 22Rv1 and C4-2B cell lines undergoing enzalutamide treatment. Notably, reducing RFX6 expression in conjunction with enzalutamide significantly decreased cell proliferation **(Extended Data Fig. 7a)**. Conversely, enhancing RFX6 expression restored proliferation in enzalutamide-treated cells **(Extended Data Fig. 7b)**, as corroborated by colony formation experiments **(Extended Data Fig. 7c-f)**. Further analysis showed that lower RFX6 levels correlated with decreased half-maximal inhibitory concentration (IC50) values for enzalutamide, indicating heightened drug sensitivity **(Extended Data Fig. 7g)**. On the other hand, elevated RFX6 expression led to higher IC50 values, suggesting reduced drug responsiveness **(Extended Data Fig. 7h)**.

Our results suggest that RFX6 could be a potential therapeutic biomarker for enzalutamide-resistant PCa. To test this, we used the C4-2B-EnzR (C4-2R) cell line, which had been cultured with enzalutamide for six months to develop drug resistance. When we treated these C4-2R cells with RFX6 shRNA, a marked decrease in cell proliferation and colony formation was observed **(Fig. 8b, c)**. In contrast, C4-2R cells treated with either enzalutamide or DMSO showed no significant difference, highlighting the potential of RFX6 downregulation to restore enzalutamide sensitivity in resistant cell lines **(Fig. 8b, c)**.

To confirm the role of RFX6 in enzalutamide-resistant PCa *in vivo*, we developed a xenograft mouse model using C4-2R cells. The enzalutamide-treated and control groups showed no significant differences in tumor volume or weight, demonstrating the resistance of cells to enzalutamide **(Fig. 8d-g)**. However, the group with RFX6 knockdown exhibited a notable reduction in both tumor volume and weight compared to the control group **(Fig. 8f-g)**. Moreover, combining RFX6 knockdown with enzalutamide treatment led to even further decreases in tumor volume and weight **(Fig. 8f-g)**. These findings indicate that depleting RFX6 not only suppresses C4-2R cell proliferation *in vitro* but also *in vivo*, highlighting the potential of RFX6 as a valuable biomarker and therapeutic target in enzalutamide-resistant PCa.

## Discussion

PCa remains a significant health challenge, and understanding its genetic and molecular underpinnings is crucial. This study uncovers the in vitro and in vivo roles and clinical relevance of RFX6 in PCa. We demonstrate that RFX6 expression is transcriptionally reprogrammed, likely due to an AR binding site gain at the PCa risk-associated rs339331/6q22 locus over human prostate tumorigenesis. We establish that RFX6 upregulates HOXA10 and activates the TGFβ pathway, promoting EMT and enhancing PCa metastasis **(Fig. 8h)**. Crucially, our findings reveal that diminishing RFX6 expression re-establishes sensitivity to enzalutamide in CRPC cells and tumors that has previously shown resistance, positioning RFX6 as a highly promising biomarker and therapeutic target for PCa treatment **(Fig. 8h)**.

A key finding of our study is the robust association between RFX6 regulatory control at the 6q22 PCa susceptibility locus and AR signaling. A nonmutational epigenetically gained binding site of AR, a vital ligand-dependent transcription factor within the steroid hormone receptor family, plays a critical role in regulating epithelial differentiation and cell proliferation, processes fundamental prostate development^33, 34, 70, 71^. This study uncovers a reprogrammed AR binding site at the 6q22 locus in PCa tumors, specifically linked to the rs339331 risk allele^31, 34^. The chromatin occupancy of AR at the rs339331 enhancer and its consequential impact on RFX6 expression shed light on the pivotal role of RFX6 in PCa progression. The notable correlation observed between RFX6 and AR expression in PCa tissues underscores the importance of their interplay in influencing tumor growth and development.

RFX6 is typically expressed in the pancreas and gastrointestinal tract, essential for pancreatic development and insulin production, with its dysregulation implicated in gastrointestinal disorders. Yet, its role in cancer is less defined. Our study reveals a novel transcriptional reprogramming of RFX6 in PCa by an interplay of disregulated genetic and epigenetic mechanisms converging on a reprogrammed binding site of the prostate lineage transcription factor AR **(Fig. 8h)**. Abnormal expression of RFX6 selectively modulates HOXA10 expression, mirroring the gene regulatory specificity seen in the activation of CHD4 by the transcription factor ZNF410^72^. The HOX gene family member HOXA10 is linked to various cancers^63, 73–76^, with its overexpression associated with adverse outcomes in PCa as described above. Our analyses also establish a strong association between RFX6 and HOXA10 expression in PCa, underscoring the critical role of HOXA10 in mediating RFX6 oncogenic impact. Furthermore, we delineate a direct regulatory axis comprising RFX6, HOXA10, and TGFβ2, wherein HOXA10 engages directly with the TGFβ2 promoter. This mechanistic understanding enhances our comprehension of RFX6 contributing to PCa cellular aggressiveness and tumor progression.

Patients with aggressive PCa often undergo ADT treatment, initially effective in extending survival^77^. However, this treatment commonly leads to CRPC, characterized by drug resistance and increased mortality^8, 33,77^. Our study reveals a novel link between RFX6 expression and resistance to enzalutamide in PCa. We found that reducing RFX6 levels enhances the sensitivity of enzalutamide-resistant PCa cells to the drug, suggesting RFX6 as a potential indicator for treatment efficacy. This finding offers new insights for therapeutic strategies in managing resistant PCa cases.

In summary, our study elucidates the multifaceted role of RFX6 in PCa, linking its regulation to epigenetically reprogrammed AR chromatin association and androgen signaling, and influencing HOXA10 gene expression. This regulatory mechanism significantly impacts the EMT and cancer metastasis and is potentially associated with enzalutamide resistance. Our insights contribute to understanding the molecular underpinnings of PCa progression and pave the way for innovative therapeutic strategies targeting RFX6^high^ expression tumors. Further study is needed to fully explore the clinical potential of RFX6 in PCa treatment and to confirm its utility as a therapeutic marker in patient management.

## Materials and Methods

### Ethics Statement

The use of human specimens in this study was approved by the Ethics Committee Board at the School of Basic Medical Sciences, Shanghai Medical College of Fudan University (approval number: 2022-Y015). All procedures involving human specimens strictly followed the ethical guidelines of the Declaration of Helsinki. Furthermore, written informed consent was obtained from each participating patient, ensuring the highest standards of ethical conduct in our medical research.

The animal experiments in this study received approval from the Animal Care and Use Committee of the School of Basic Medical Sciences at Shanghai Medical College of Fudan University (approval number: 20220228-014). These protocols complied rigorously with the National Institutes of Health Guide for the Care and Use of Laboratory Animals. This adherence guarantees the ethical and humane treatment of all animals involved in our research.

### Animal experiments

For the xenograft mouse model, either 22Rv1 or C4-2R stable cell lines (5 × 10^6^ cells) were mixed with Matrigel at a 1:1 ratio and subcutaneously injected into the right rear flank of 6-week-old male nude mice. Tumor size and volume were monitored at three-day intervals and calculated using the formula: volume = length × (width)^2^/2. When tumors reached a volume of 50-100 mm^3^, mice in the enzalutamide group received oral doses of 10 mg/kg enzalutamide. Approximately 30 days post-injection, mice were euthanized, and tumors were harvested for hematoxylin and eosin staining and immunohistochemistry analysis.

For the lung metastasis model, 2 × 10^6^ fluorescein-labeled 22Rv1 cells were suspended in 100 μL PBS and injected into the tail vein of SCID mice. Immediately post-injection, D-luciferin potassium salt (Yeasen) was administered intraperitoneally, followed by *in vivo* imaging to confirm successful injection. Starting from week 4, *in vivo* imaging was conducted weekly. All animal experiments were performed following ethical guidelines approved by the Department of Laboratory Animal Science at Fudan University.

### Cell culture

All cell lines were acquired from the American Type Culture Collection (ATCC) and were routinely tested for mycoplasma contamination. None of these cell lines were found to be contaminated with mycoplasma during the course of our study. VCaP, DU145, and 293T cells were cultured in DMEM (Thermo Fisher Scientific), while 22Rv1, PC3, and C4-2R cells were cultured in RPMI1640 (Thermo Fisher Scientific). All culture media were supplemented with antibiotics (penicillin and streptomycin, Meilunbio) and 10% Fetal Bovine Serum (FBS, Gibco). For experiments involving AR activity, androgen-responsive PCa cells were treated with 100 nM dihydrotestosterone (DHT; Sigma) for specified durations.

### Lentiviral constructs, lentivirus production and infection

To establish RFX6 and HOXA10 overexpression cell lines, we obtained their gene coding sequences (CDS) from NCBI and designed primers. These primers, synthesized by Sangon Biotech, amplified the CDS sequences from PCa cell line cDNA. The amplified sequences, along with corresponding epitope tag fragments, were inserted into the pLVX-EF1a vector. For shRNA constructs, we used the pLKO.1-puro vector, selecting the two most efficient shRNAs targeting each gene from a set of six (details in **Supplementary Table 2**). Lentiviral particles expressing shRNAs were produced using a third-generation packaging system in HEK 293T cells (ATCC, CRL-11268). At 70-80% confluence, 293T cells in 6 cm plates were co-transfected with the lentiviral transfer vector, envelope plasmid pVSVG, packaging plasmids pRSV-Rev and pMDLg/pRRE, using PEI. The culture medium was replaced after 6-8 hours, and viral supernatant was collected at 48-hour intervals for 4 days. This supernatant, containing viral particles, was filtered and either stored at -80 °C or used immediately for cell infection. Target cells at 70-80% confluence were infected with the viral supernatant supplemented with 8 μg/ml polybrene (Sigma: TR-1003-G). Infected cells were selected using puromycin (Meilunbio) - 1 μg/ml for 22Rv1, C4-2B, and C4-2R cells, and 0.5 μg/ml for DU145 and PC3 cells - added 36 to 48 hours post-infection. After 3 days of screening, the surviving cells were split and maintained at the same puromycin concentration.

### Cell proliferation experiment

For viability assessment, 22Rv1, C4-2B, or C4-2R cells were seeded at a density of 4×10^3^ cells/well), and PC3 or Du145 cells at 1×10^3^ cells/well, in 96-well plates. Post-adhesion, 22Rv1 cells were treated with 0.5 mg/ml MTT (Beyotime) for 4 hours. After removing the supernatant, DMSO was used to dissolve the formazan crystals. The optical density (OD) was measured at a wavelength of 490 nm. Other cell lines were incubated with CCK8 (Beyotime, Cell Counting Kit-8) for 1 hour, with OD readings taken at 450 nm. Subsequent OD measurements were taken every 48 hours. Statistical analyses were performed after 3-4 time points.

### Colony formation assay

22Rv1, C4-2B or C4-2R cells (1×10^3^ cells/well), PC-3 or DU145 cells (500 cells/well) were seeded in 12-well plates. After 2 weeks of growth, cells were fixed with 4% paraformaldehyde and stained with 0.05% crystal violet.

### Cell migration and invasion assays

Stable cell lines were resuspended in DMEM or RPMI-1640 medium containing 0.5% FBS to archieve a concentration of 2.5 × 10^5^ cells/ml. For each assay, 0.2 ml of the cell suspension was seeded into 8-μm Transwell inserts (Corning Costar). Inserts were either coated with Matrigel (Yeasen) for invasion assays or uncoated for migration assays. The lower chambers of the wells were filled with 0.5 ml of RPMI-1640 supplemented with 10% FBS. After 48 hours of incubation, cells remaining on the upper surface of the membranes were gently removed. The membranes were then fixed and stained with crystal violet (Sangon Biotech) for 6 hours. Cells that had migrated or invaded through the membrane were quantified by counting them in eight different microscopic fields per membrane, using a 10× magnification). Data were collected from five replicate inserts per condition. Statistical analyses of the results were performed using a two-tailed t test.

### Cell apoptosis assay

Cells were seeded in a 24-well plate at a density of 5 × 10^4^ cells per well. Each condition was set up in triplicate. After 24 hours of incubation, cells were harvested and processed as per the instructions provided by Annexin V-FITC Apoptosis Detection Kit (Beyotime). Immediately following processing, the cells were analyzed using flow cytometry to assess apoptosis.

### Quantitative RT-PCR

RNA samples were extracted from cultured cell lines using the EZ-10 DNAaway RNA Mini-Preps Kit (Sangon Biotech: B618133), following the manufacturer’s instructions. The isolated RNA was reverse transcribed into cDNA using the PrimeScript™ RT Master Mix (Takara: RR036B) according to the the protocol provided by the manufacturer. Quantitative RT-PCR analysis was performed using ChamQ universal SYBR qPCR master mix (Vazyme: Q711-02), according to the product user guides. The RT-PCR was conducted on a LightCycler 480 Instrument II (Roche). Primer pairs for each target gene were meticulously designed and evaluated to ensure specificity and efficiency. Only those demonstrating high specificity and amplification efficiency were utilized for RT-PCR quantification. The sequences of the primer used in this study are listed in **Supplementary Table 3**.

### Western Blot and antibody

Cells were first washed with phosphate-buffered saline (PBS) and then lysed using Nonidet P-40 lysis buffer containing 1% protease inhibitor and 1% phosphatase inhibitor. The lysis was carried out at 4 °C for 30 minutes. The cell lysates were then mixed with sodium dodecyl sulphate–polyacrylamide gel electrophoresis (SDS-PAGE) sample loading buffer and heated at 95 °C for 10 minutes. Following heating, the samples were subjected to SDS-PAGE followed by Western blot analysis for protein expression detection. Detailed information about all the antibodies used in this study can be found in **Supplementary Table 4**.

### Luciferase reporter assay

In the luciferase activity assay, we initially seeded 5 × 10^3^ 22Rv1 cells per well in triplicate in a 96-well plate and allowed them to incubate for 24 hours. Transfection was then conducted using Lipofectamine 3000 (Invitrogen), following the manufacturer’s instructions. For each well, the transfection mixture consisted of 100 ng of PGL4.10 luciferase reporter plasmid, 100 ng of a transcription factor plasmid, and 2 ng of the PGL4.75 (Renilla luciferase) plasmid. After a 48-hour transfection period, we quantified the luciferase and Renilla signals using the dual-luciferase reporter gene assay kit (Promega, E1960), following the protocol provided by the manufacturer.

### Chromatin immunoprecipitation (ChIP)

Cells were cross-linked with 1% formaldehyde at room temperature for 10 minutes, followed by the addition of 125 mM glycine to quench the reaction. Subsequently, cell pellets were harvested and resuspended in hypotonic lysis buffer (10 mM KCl, 20 mM Tris–HCl pH8.0, 10% Glycerol, 2 mM DTT), and gently rotated at 4 °C for 50 minutes to isolate nuclei. Nuclei were then washed twice with cold PBS and resuspended in SDS lysis buffer (50 mM Tris– HCl, pH8.1, 10 mM EDTA, 0.5% SDS, 1×Protease inhibitor cocktail (Bimake)). Chromatin was sheared to ∼250 bp fragments using a M220 Focused-ultrasonicator (Covaris). For immunoprecipitation, 70 μl of Dynabeads Protein G slurry (Thermo Fisher Scientific) was prepared by washing twice with blocking buffer (0.5% BSA in IP buffer) and added to each sample, followed by incubation with 6 μg of either anti-HOXA10 antibody or control IgG at 4 °C for 12 hours. The supernatant was then removed. For chromatin binding, 250 μg soluble chromatin in IP buffer (20 mM Tris–HCl pH8.0, with 2 mM EDTA, 150 mM NaCl, 1%Triton X-100, and Protease inhibitor cocktail) was added to the bead-antibody complexes and incubated for another 12 hours. The supernatant was then removed and the bead-DNA–protein complexes was washed six times with RIPA washing buffer (50 mM HEPES pH 7.6, 1 mM EDTA, 0.7% sodium deoxycholate, 1% NP-40, 0.5 M LiCl). Subsequently, the DNA– protein complexes were eluted in DNA extraction buffer (10 mM Tris–HCl pH 8.0, 1 mM EDTA, and 1% SDS), followed by overnight treatment with Proteinase K and RNase A at 65 °C to reverse cross-linking. The immunoprecipitated DNA was purified using QIAquick PCR or Mini-Elute PCR purification kits (Qiagen) and analyzed by massive parallel sequencing or quantitative real-time PCR assay.

### RNA-seq processing

The quality of RNA-seq data was evaluated using FastQC (version: 0.11.9) (www.bioinformatics.babraham.ac.uk/projects/fastqc/). Low-quality reads were trimmed, and adapters were removed utilizing AdapterRemoval (version: 2.3.2)^78^. The cleaned data were then mapped to the reference genome (hg38) using STAR (version: 2.7.9a)^79^. The aligned BAM files were sorted with SAMtools (version: 1.13)^80^. Gene read counts were derived from the sorted BAM files using HTseq-count (version: 0.13.5)^81^. DESeq2^82^ package was used for identifying differentially expressed genes (DEGs, applying a threshold of |Fold Change| > 2 and False Discovery Rate (FDR) < 0.05).

### Gene expression correlation analysis

We performed the co-expression analysis to evaluate the expression correlation between RFX6 and AR or HOXA10 from multiple independent PCa cohorts. We employed both Pearson’s product-moment correlation and Spearman’s rank correlation rho methods for all linear expression correlation tests. This dual approach allowed for a comprehensive evaluation of the relationships between these genes in the context of PCa.

### Survival analysis

We employed Kaplan-Meier survival analysis to evaluate the prognostic impact of RFX6 or HOXA10 on PCa outcomes across multiple independent cohorts. Patients were first stratified into two groups based on their mean expression levels of the genes of interest. To assess the synergistic effect of rs339331 genotype and RFX6 expression on PCa patient survival, we first stratified patients by the rs339331 genotype. Subsequently, we analyzed the survival implications of RFX6 expression in patients with TT or TC/CC alleles. The Kaplan-Meier survival curves were generated using the R package ‘Survival’ (v. 3.2.3), with differences between curves evaluated using log-rank test.

### Statistics

Statistical analysis was performed using GraphPad Prism 9 software. Bar graphs data are presented as fold change or percentage relative to the control, with standard deviations (SD) derived from three independent experiments. The student’s t test was employed for the analysis of normally distributed data. The log-rank test was used for the patient survival analysis. A P-value < 0.05 was considered statistically significant. Levels of significance were indicated as follows: *P < 0.05, **P < 0.01, ***P < 0.001, and ****P < 0.0001. Statistical tests for investigating RFX6 or HOXA10 expression levels across normal tissues, primary PCa tumor, and metastatic tissues, as well as clinical features like PSA, and tumor stage, were analyzed using the Mann-Whitney U test or the Kruskal-Wallis H test, based on the number of comparison groups. For the results from microarray-based expression profiling, gene probes with lowest P values were selected. Samples with missing gene expression or patient survival data were excluded from analyses. RStudio (version 1.4.1106) with R (version v. 4.1.0) was used for statistical testing.

### Data availability

In this study, we have utilized several public datasets, which include GSE3325, GSE3868, GSE6811, GSE35988, GSE6099, GSE70770, GSE2109, GSE21034, GSE21032, GSE62872 from GEO database (https://www.ncbi.nlm.nih.gov/geo/), CPGEA (http://bigd.big.ac.cn/gsa-human/, accession number is PRJCA001124), GTEx (https://gtexportal.org/home/), TCGA (URLs: TCGA data matrix, https://tcga-data.nci.nih.gov/tcga/dataAccessMatrix.htm; TCGA Research Network, http://cancergenome.nih.gov/), and MSKCC (https://www.mskcc.org/). In addition, our study utilized public cohort data from Stockholm, FHCRC, MCTP, Tmm, DKFZ and SU2C that can be accessed via the cBioPortal (https://www.cbioportal.org/). All types of data supporting the discoveries in this study are available upon reasonable request from the corresponding authors.

## Acknowledgements

We express our sincere gratitude to the members of the Wei laboratory for their valuable discussions and insightful comments, which significantly contributed to the improvement of this work. Our research was generously supported by several funding sources: The National Natural Science Foundation of China (82073082; 82311530050), the Shanghai Interactional Collaborative Project (23410713300), the National Key Research and Development Program of China (2022YFC2703600), Jane ja Aatos Erkon säätiö, Sigrid Juséliuksen Säätiö, Syöpäjärjestöt, and the Fudan recruit funds. We acknowledge that the funding agencies played no role in the design, execution, analysis, or interpretation of the study, nor in the writing and publication of this work. We also extend our thanks for the Linux High-throughput Computing servers provided by the Medical Research Data Center in Shanghai Medical College of Fudan University, and the CSC-IT CENTER FOR SCIENCE LTD.

## Author contributions

M.Z. and G.-H.W. designed the project and analyzed the majority of the experiments. G.-H.W. conceived the study and provided supervision, suggestions, and overall guidance. M.Z. performed most of the experiments with assistance from P.T., L.L., Q.Z., and Y.L. W.X., Q.Z., and Z.W. were responsible for the bioinformatics analysis and clinical evidence-based data mining. M.Z. and G.-H.W. wrote the manuscript. All authors actively participated in discussing the results and contributing to the manuscript.

