## Extended Data for "RFX6 at locus 6q22 confers metastasis and drug resistance in prostate cancer"

**Zhong *et al*. Extended Data Figure 1**

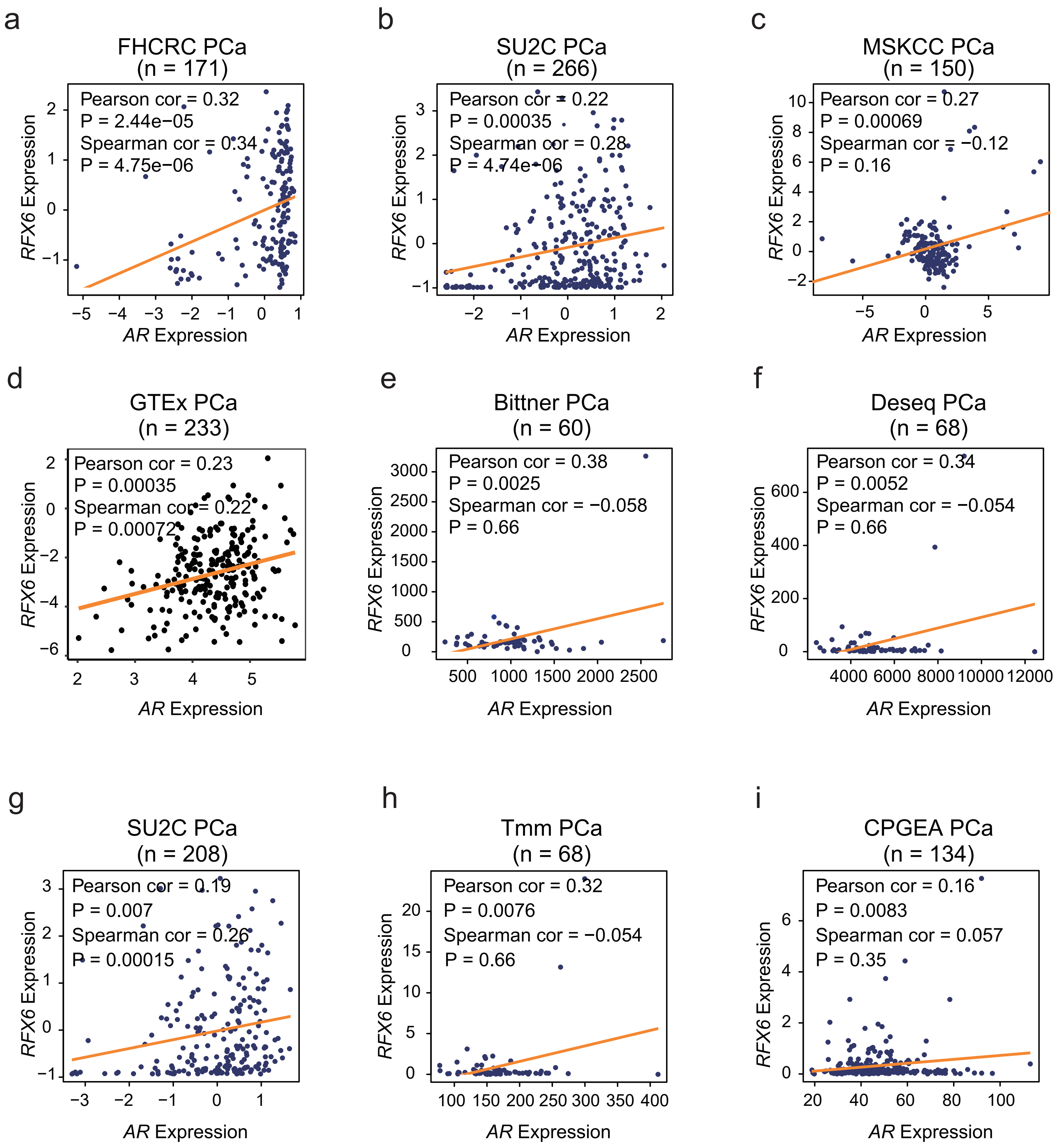

**Extended Data Figure 1. Strong correlation between RFX6 and AR expression in PCa.**

**a-i**, catter plots demonstrating a positive correlation between RFX6 and AR expression across various PCa patient cohorts, including FHCRC PCa (n = 171), SU2C PCa (n = 266, 208), MSKCC PCa (n = 150), GTEx PCa (n = 233), Bittner PCa (n = 60), Deseq PCa (n = 68), Tmm PCa (n = 68) and CPGEA PCa (n = 134) cohorts.

Statistical Analysis: Pearson and Spearman correlation analyses were employed to assess p-values across all cohorts.

Zhong *et al*. Extended Data Figure 2

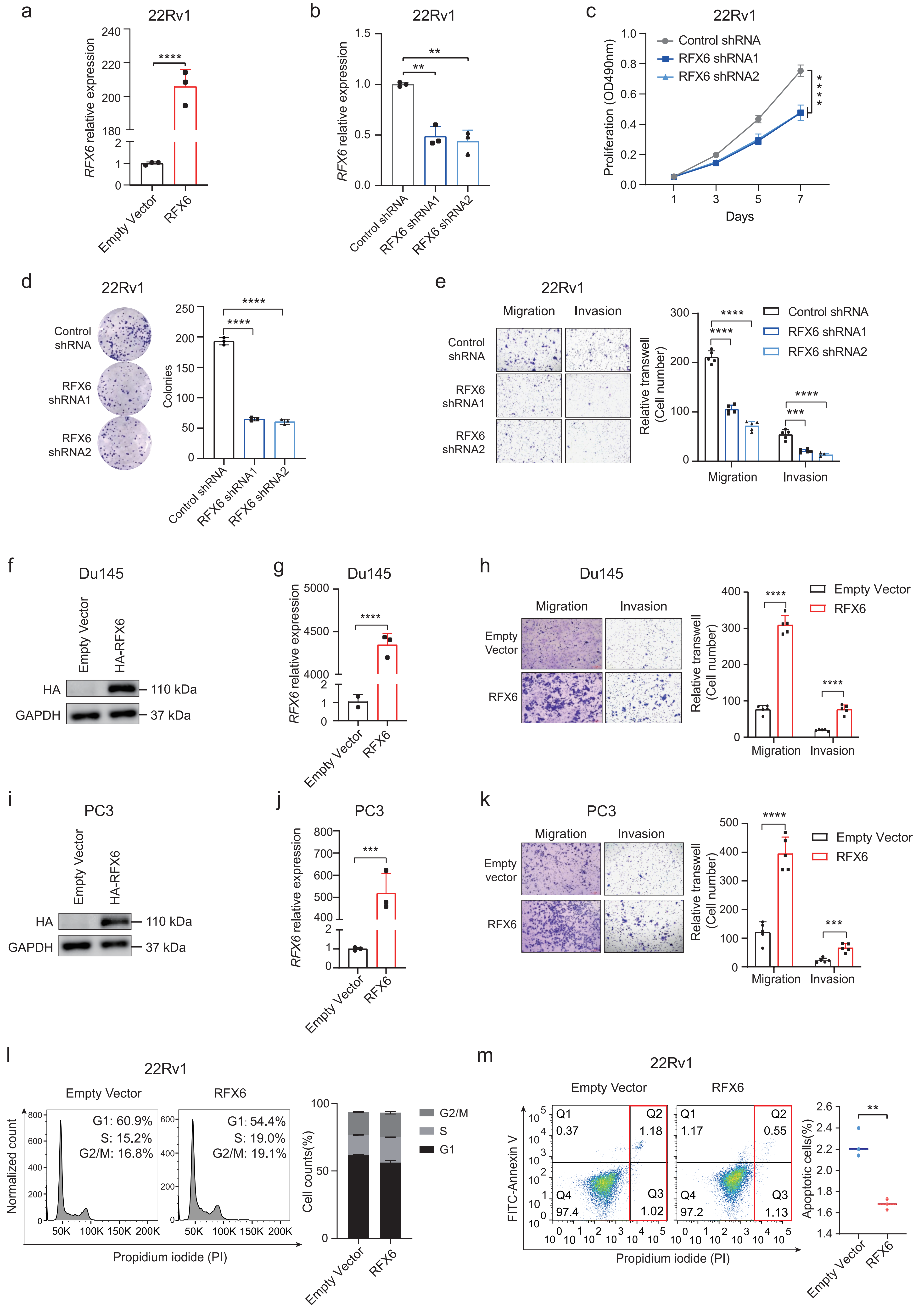

**Extended Data Figure 2. Evaluating the effects of RFX6 expression alterations on PCa cell phenotype.**

**a-b**, Analysis of RFX6 expression levels in 22Rv1 cells under various treatments, determined by quantitative real-time PCR.

**c-d**, Assessment of cellular proliferation in 22Rv1 cells with RFX6 knockdown using MTT assay ($OD_{490}$; mean ± SD from three independent experiments) (**c**) and colony formation assay (**d**).

**e**, Representative images and quantification of cell migration and invasion 22Rv1 cells stably expressing shRNAs targeting RFX6.

**f**, **i**, Immunoblots showing RFX6 protein levels in Du145 (**f**) and PC3 (**i**) cells with either empty vector or RFX6 overexpression.

**g**, **j**, Real-time PCR analysis of RFX6 mRNA levels normalized to GAPDH in DU145 (**g**) and PC3 (**j**) cells with varying RFX6 expression.

**h**, **k**, Representative images and quantification of cell migration and invasion in DU145 (**h**) or PC3 (**k**) cells with the specified RFX6 expression status.

**l**, Cell cycle distribution analysis of 22Rv1 cells with empty vector or RFX6 overexpression, conducted via flow cytometry, focusing on the G1, S, and G2/M phases.

**m**, Apoptosis assessment in 22Rv1 cells with empty vector or RFX6 overexpression using flow cytometry.

All the statistical significance assessed using the two-tailed Student's t tests. Significance levels indicated as *$p < 0.05$, **$p < 0.01$, ***$p < 0.001$, ****$p < 0.0001$.

**Zhong *et al*. Extended Data Figure 3**

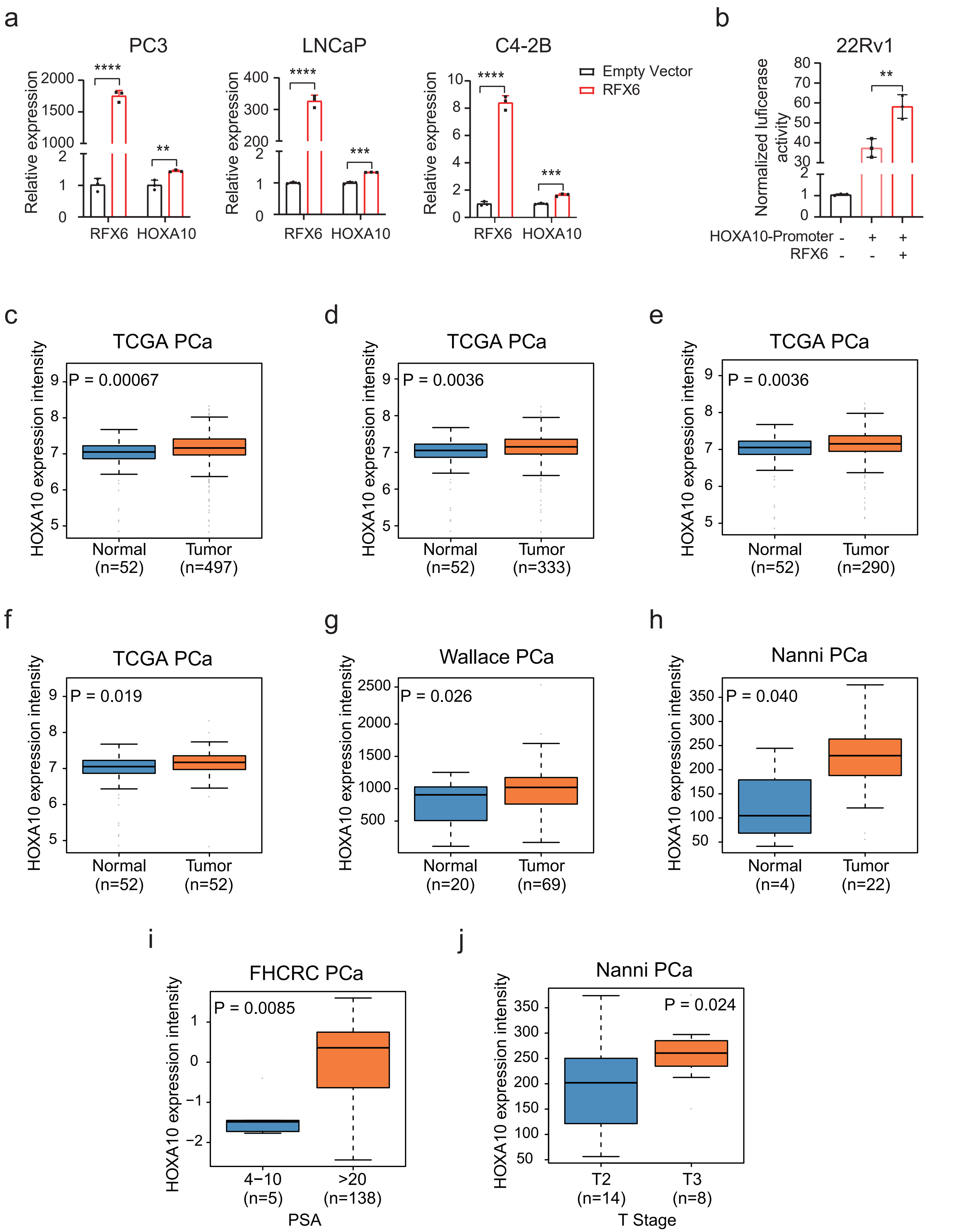

**Extended Data Figure 3. RFX6 regulation of HOXA10 expression in various PCa cell lines.**

**a**, Analysis of HOXA10 expression normalized to GAPDH in PC3, LNCaP, and C4-2B cell lines with stable RFX6 overexpression, measured by quantitative real-time PCR.

**b**, Luciferase reporter assay demonstrating enhanced transcriptional activity of the HOXA10 promoter in 22Rv1 cells with RFX6 overexpression compared to empty vector control.

**c-h**, Clinical data analyses indicating a correlation between HOXA10 upregulation and human prostate tumor progression.

**i-j**, Clinical data showing associations of HOXA10 upregulation with elevated levels of prostate-specific antigen (PSA) (**i**) and higher tumor stage (**j**) in PCa.

Statistical Analysis: Two-tailed Student's t-tests were employed for all data points to assess statistical significance and p-values, with significance levels denoted as $*p < 0.05$, $**p < 0.01$, $***p < 0.001$, $****p < 0.0001$.

**Zhong *et al*. Extended Data Figure 4**

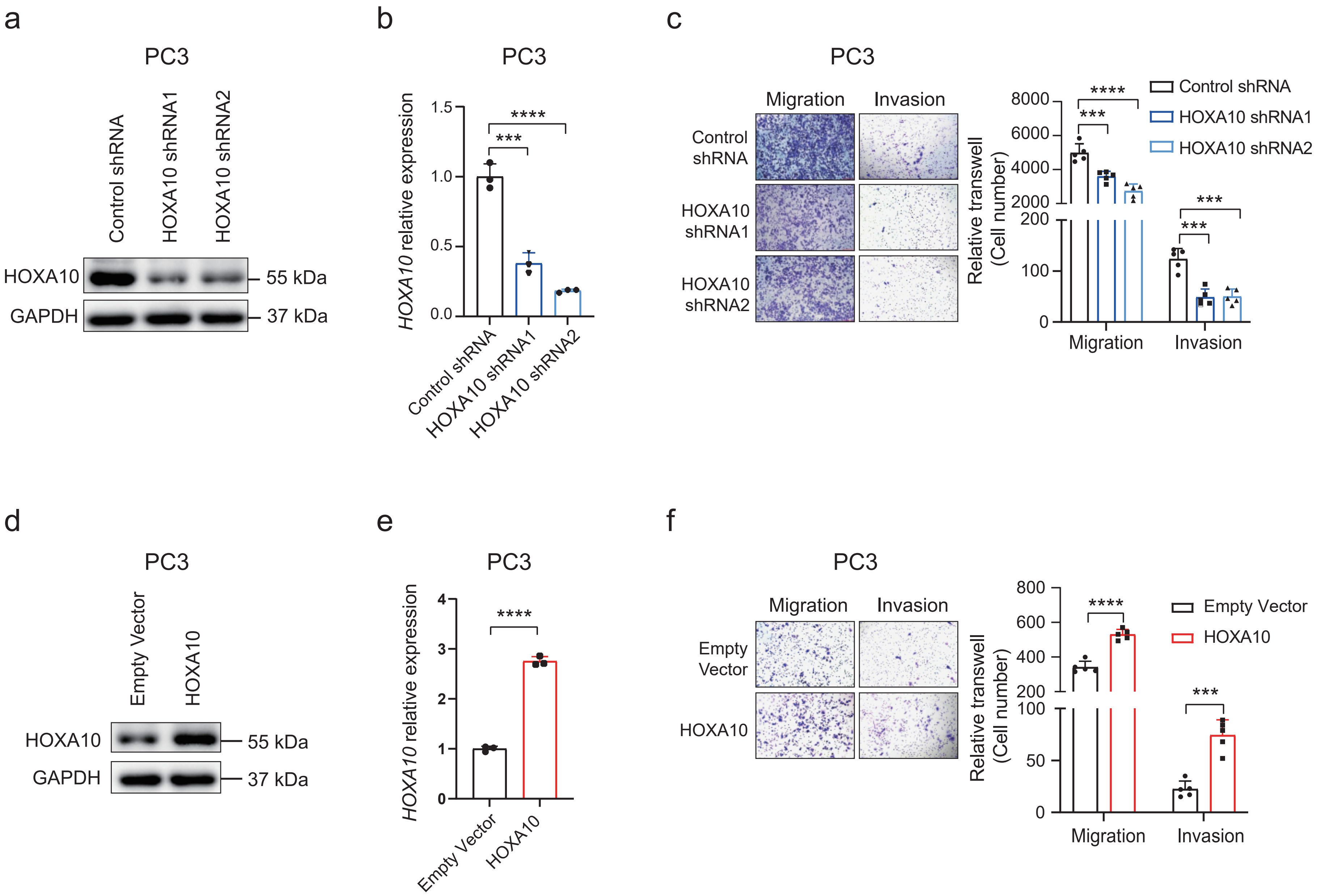

**Extended Data Figure 4. Investigating the effects of HOXA10 expression alterations on PC3 cell phenotype.**

**a-b**, Verification of shRNA-mediated RFX6 knockdown efficiency in PC3 cells, assessed by immunoblotting (**a**) and quantitative real-time PCR (**b**).

**c**, Representative images showcasing migration and invasion of PC3 cells stably expressing shRNAs targeting RFX6.

**d**, Immunoblots illustrating HOXA10 protein levels in PC3 cells with either empty vector or HOXA10 overexpression.

**e**, Quantitative real-time PCR analysis of relative HOXA10 mRNA expression levels normalized to GAPDH in PC3 cells with empty vector or HOXA10 overexpression.

**f**, Representative images of migration and invasion assays in PC3 cells transfected with either empty vector or HOXA10 expression construct.

Statistical Analysis: All data were analyzed for statistical significance using two-tailed Student's t-tests. Significance levels are indicated as $*p < 0.05$, $**p < 0.01$, $***p < 0.001$, $****p < 0.0001$.

**Zhong *et al*. Extended Data Figure 5**

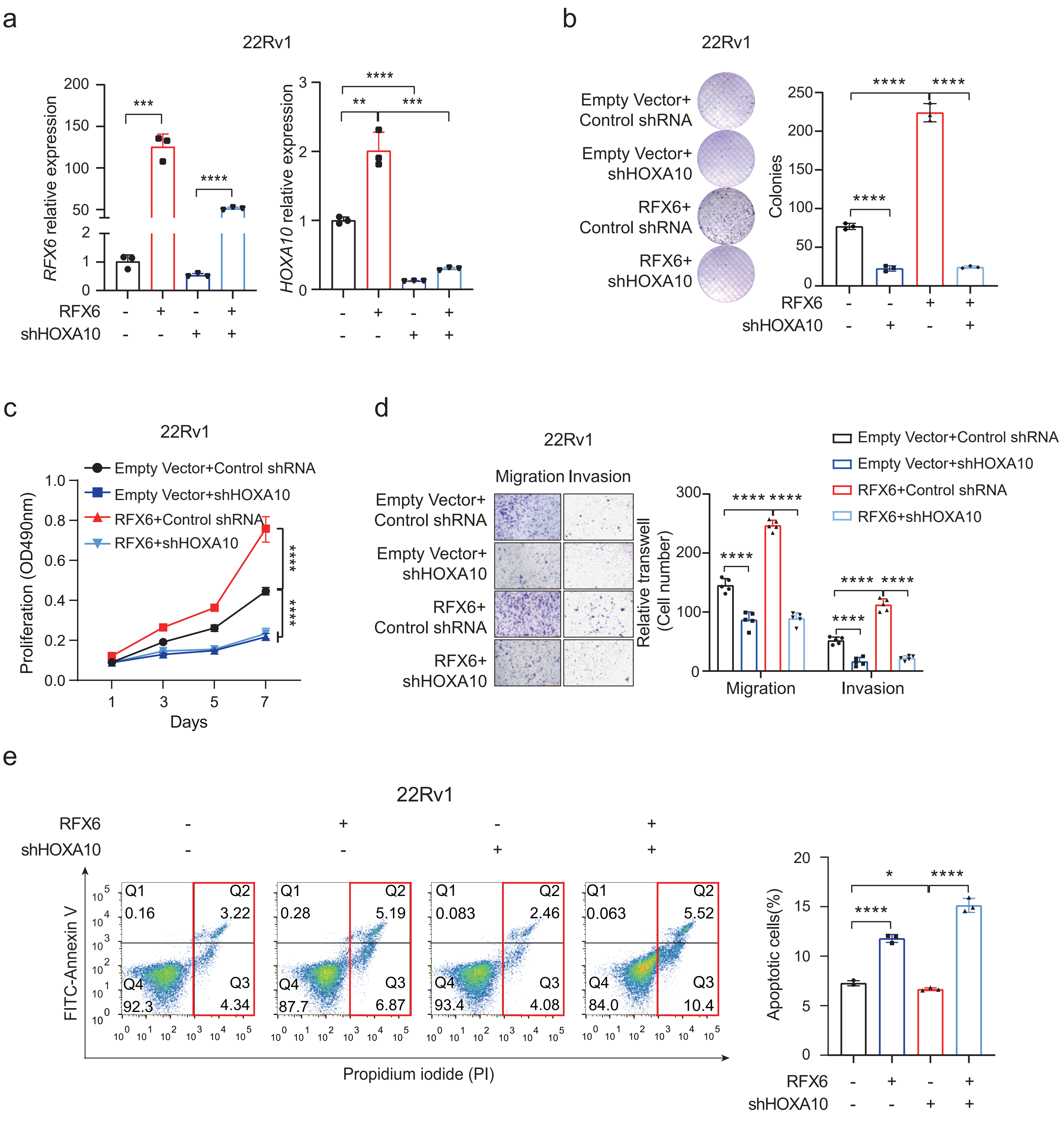

**Extended Data Figure 5. Reversal of RFX6-induced cellular phenotype by HOXA10 suppression *in vitro*.**

**a**, Analysis of relative expression levels of RFX6 and HOXA10 in 22Rv1 cells under various treatments, measured by quantitative real-time PCR.

**b-c**, Evaluation of cellular proliferation in 22Rv1 cells with the specified treatments using colony formation assay (**b**) and MTT assay ($OD_{490}$; mean ± SD from three independent experiments) (**c**).

**d**, Representative images and analysis of cell migration and invasion in 22Rv1 cells stably expressing different treatment groups.

**e**, Flow cytometry analysis to determine the proportion of apoptotic cells in the indicated treatment groups.

Statistical Analysis: Statistical significance for all data points was determined using two-tailed Student’s t-tests. Significance levels are denoted as *p < 0.05, **p < 0.01, ***p < 0.001, ****p < 0.0001.

Zhong *et al*. Extended Data Figure 6

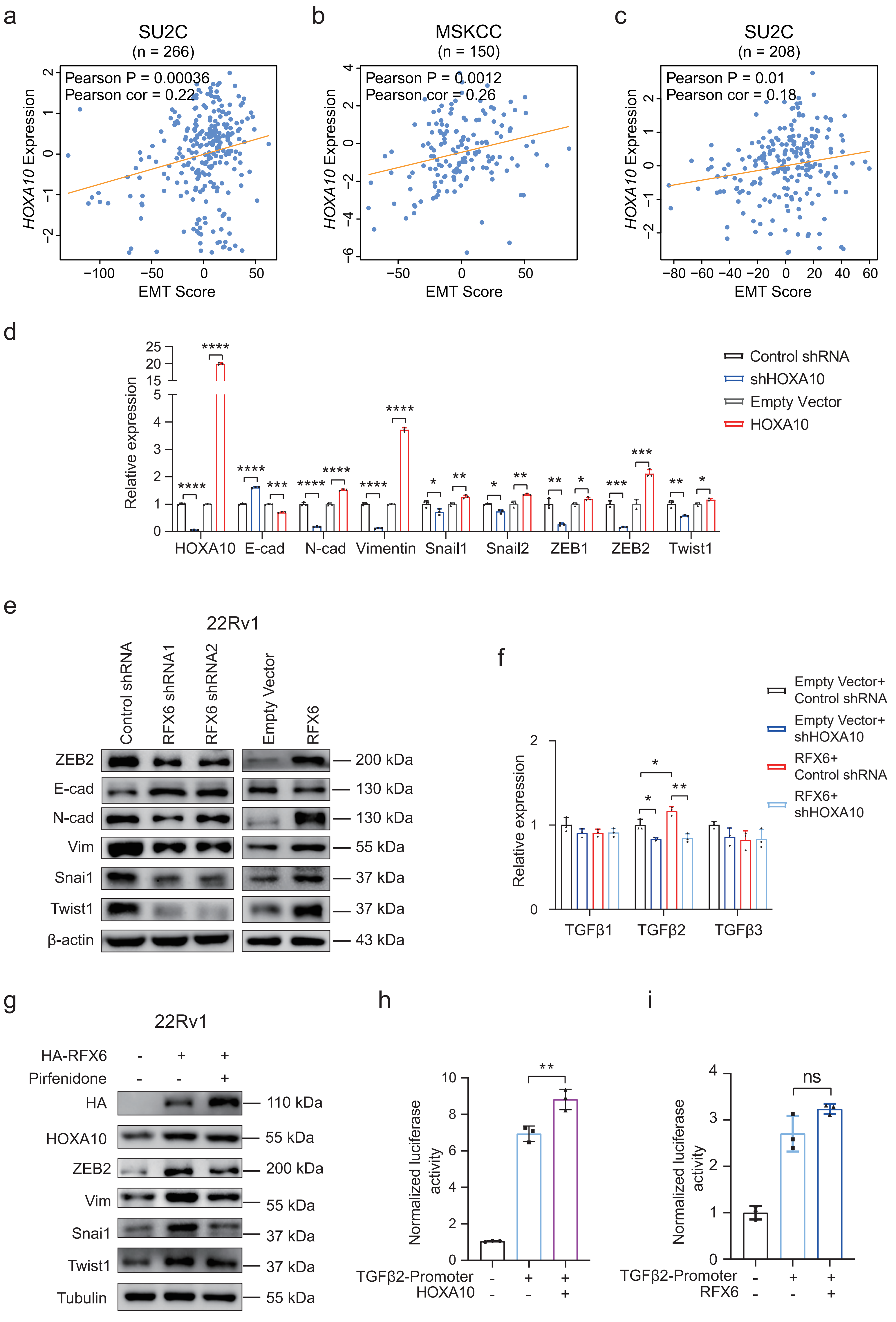

**Extended Data Figure 6. RFX6 and HOXA10 expression impact on EMT marker expression via TGFβ2 modulation.**

**a-c**, Positive correlation between HOXA10 expression and computed EMT scores in various independent PCa cohorts.

**d**, Real-time PCR analysis of EMT-related molecules (E-cadherin, N-cadherin, Vimentin, Snail, Slug, ZEB1, ZEB2, Twist1) in cells with specified treatments.

**e**, Immunoblotting determination of EMT-related molecules (E-cadherin, N-cadherin, Vimentin, Snail1, ZEB2, Twist1) in cells under indicated treatment conditions.

**f**, Quantitative analysis of TGFβ1, TGFβ2, and TGFβ3 mRNA levels in 22Rv1 cells with various treatments.

**g**, Assessment of EMT-associated molecules (E-cadherin, N-cadherin, vimentin, Snail, ZEB2, Twist1) in 22Rv1 cells following Pirfenidone treatment, using immunoblotting.

**h-i**, Luciferase reporter assays evaluating the impact of HOXA10 (**h**) and RFX6 (**i**) on the transcriptional activity of the TGFβ2 promoter in 22Rv1 cells, compared to the empty vector control.

Statistical Analysis: All data were analyzed for statistical significance using two-tailed Student's t-tests. Significance levels are indicated as *$p < 0.05$, **$p < 0.01$, ***$p < 0.001$, ****$p < 0.0001$.

**Zhong *et al*. Extended Data Figure 7**

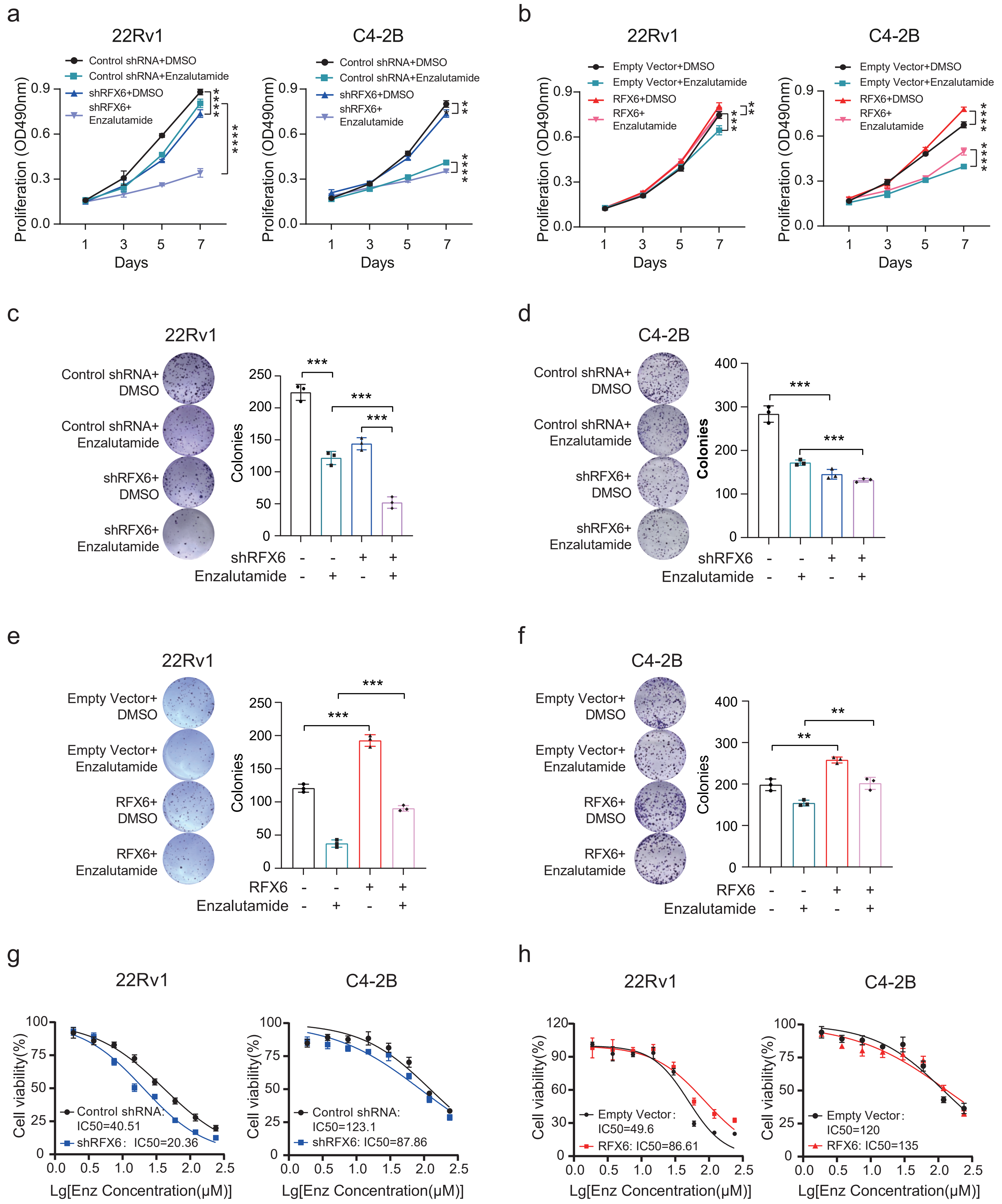

**Extended Data Figure 7. Effects of RFX6 levels on enzalutamide sensitivity in CRPC cell lines.**

**a-f**, Evaluation of cellular proliferation in 22Rv1 or C4-2B cells under various treatments, measured using MTT assay ($OD_{490}$; mean ± SD from three independent experiments) (**a-b**) and colony formation assay (**c-f**).

**g-h**, Determination of the half-maximal inhibitory concentration (IC50) of enzalutamide in 22Rv1 or C4-2B cells, based on the indicated treatments.

Statistical Analysis: Two-tailed Student's t-tests were utilized to assess statistical significance across all data points. Significance levels are denoted as $^*p < 0.05$, $^{**}p < 0.01$, $^{***}p < 0.001$, $^{****}p < 0.0001$.
